# Multiple timescales of neural dynamics and integration of task-relevant signals across cortex

**DOI:** 10.1101/2020.02.18.955427

**Authors:** Mehran Spitmaan, Hyojung Seo, Daeyeol Lee, Alireza Soltani

**Author notes:** Corresponding author: AS, Department of Psychological and Brain Sciences, Dartmouth College, Hanover NH 03755,.

## Abstract

Recent studies have proposed the orderly progression in the time constants of neural dynamics as an organizational principle of cortical computations. However, relationships between these timescales and their dependence on response properties of individual neurons are unknown. We developed a comprehensive method to simultaneously estimate multiple timescales in neuronal dynamics and integration of task-relevant signals along with selectivity to those signals. We found that most neurons exhibited multiple timescales in their response, which consistently increased from parietal to prefrontal to cingulate cortex. While predicting rates of behavioral adjustments, these timescales were not correlated across individual neurons in any cortical area, resulting in independent parallel hierarchies of timescales. Additionally, none of these timescales depended on selectivity to task-relevant signals. Our results not only suggest multiple canonical mechanisms for increasing timescales of neural dynamics across cortex but also point to additional mechanisms that allow decorrelation of these timescales to enable more flexibility.

## Introduction

Despite the tremendous heterogeneity in terms of cell types, connectivity, and neural response across brain areas, neuroscientists have long entertained various ideas about parsimonious organizational principles that could underlie such heterogeneity (Lennie, 1998; Shepherd et al., 2005), as well as how this heterogeneity contributes to brain computations (Shamir and Sompolinsky, 2006; Goris et al., 2015). Recent studies have illustrated that the heterogeneity of neural response across the brain is anatomically ordered (Hasson et al., 2008; Honey et al., 2012; Goris et al., 2014; Meder et al., 2017; Murray et al., 2014). For example, intrinsic timescales of neural fluctuations, presumably reflecting circuit dynamics, increase from sensory to prefrontal cortices (Murray et al., 2014). Moreover, time constants of modulations by reward feedback (reward memory) also increase in tandem with intrinsic timescales across cortical areas (Bernacchia et al., 2011; Murray et al., 2014).

These parallel hierarchies of timescales in intrinsic fluctuations and reward memory, however, were estimated with different methods. More specifically, intrinsic timescales were estimated using the decay rate of autocorrelation in neural response across the population of neurons in a given area (Murray et al., 2014), whereas reward-memory timescales were obtained using activity profiles of individual neurons across multiple trials (Bernacchia et al., 2011). Therefore, it is unclear whether the presumed relationship between intrinsic and reward-memory timescales holds at the level of individual neurons. If these timescales are correlated across individual neurons, it would suggest that the source of intrinsic fluctuation might also underlie the persistence of task-related signals. By contrast, the absence of such a relationship among individual neurons could indicate that there are separate mechanisms underlying these timescales.

A few recent studies have shown that intrinsic timescales during the fixation period–– presumably before strong task-relevant signals emerge in the cortical activity ––can predict encoding of task-relevant signals later in the trial for some but not all cortical areas. This includes the encoding of chosen value during value-guided decision making (Cavanagh et al., 2016), persistent activity during working-memory tasks (Nishida et al., 2014; Cavanagh et al., 2018; Wasmuht et al., 2018), and activity related to upcoming behavioral responses during a visually-cued strategy task (Fascianelli et al., 2019; Cirillo et al., 2018). However, in all these studies, the decay rate of autocorrelation in the firing response of individual neurons during one epoch of the task (fixation period) is compared with encoding of task-relevant signals in other epochs of the task. This leaves open the possibility that the observed relationship might be spurious, because the same dynamic process related to intrinsic timescales might also influence the time course of task-relevant signals.

It is also unknown whether the observed hierarchies of reward-memory timescales depend on the selectivity of individual neurons relative to external or task-relevant signals (response selectivity). For example, long reward-memory timescales might also require strong reward selectivity. If so, heterogeneity in response selectivity might decorrelate reward-memory timescales from intrinsic timescales across different neurons, even if they were generated via a single mechanism. By contrast, independence of timescales and response selectivity could indicate that reward-memory and intrinsic timescales might be generated via separate mechanisms. This would challenge the idea that the hierarchies of timescales occur due to similar processing of information across multiple brain areas (Hunt and Hayden, 2017; Yoo and Hayden, 2018) and instead points to the importance of the heterogeneity of local circuits (Chaudhuri et al., 2015).

To address aforementioned questions, we developed a general and robust method to fit individual neurons’ response to estimate four distinct timescales in the activity of individual neurons along with their selectivity to task-relevant signals simultaneously (**Figure 1a–c**). We applied this method to recordings from 866 single neurons in four cortical areas across six monkeys performing the same competitive game of matching pennies (Barraclough et al., 2004).

**Figure 1.**
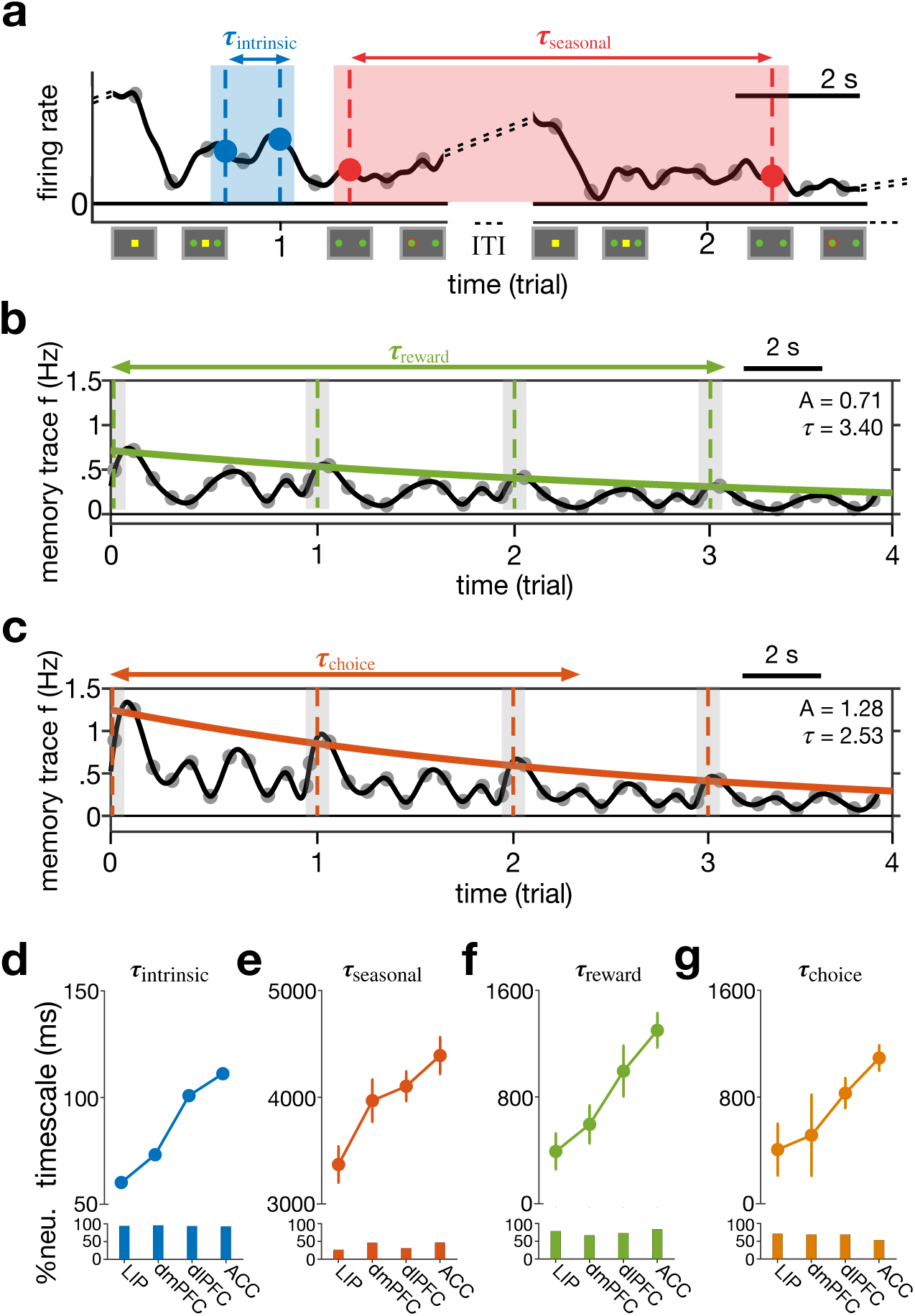
Parallel hierarchies of timescales of neural fluctuations and integration of task-relevant signals across cortex. **(a**–**c)** Simultaneous estimation of four types of timescales in neural response, illustrated for activity of an example ACC neuron. Activity in a given time epoch is related to response during previous epochs in the same trial (intrinsic timescale: *τ*_*intrinsic*_; **a**), response during the same epoch in the preceding trials (seasonal timescale: *τ*_*seasonal*_; **a**), reward outcome on previous trials measured by a memory-trace filter (reward-memory timescale: *τ*_*reward*-_; **b**), and monkeys’ choice (left vs. right) in the preceding trials (choice-memory timescale: *τ*_*choice*_; **c**). **(d–g)** Hierarchies of the four types of timescales across the cortex. Plots show the median of intrinsic (**d**), seasonal (**e**), reward-memory (**f**), and choice-memory (**g**) timescales in four cortical areas estimated using the best model for fitting response of each neuron. Error bars indicate s.e.m. Only neurons that showed significant timescales are included in each panel. Bar graphs show the fraction of neurons with a significant timescale in each area.

## Results

### General method for estimation of multiple timescales

We aimed to keep our method for estimating timescales as general as possible while capturing heterogeneity in neural response (**Figure 1a–c**). More specifically, based on previous studies (Murray et al., 2014; Bernacchia et al., 2011), we assumed that neural response at any time point in a trial could depend on activity during earlier epochs in the same trial and similar epochs in the preceding trials, as well as on reward outcome (reward vs. no reward) and choice (left vs. right) on the preceding trials. The first type of dependence—activity from epochs in the same trial—was captured by an autoregressive (AR) component that predicts spikes in a given 50-msec time bin based on spikes in the preceding 8 time bins. The autoregression coefficient for each term of this AR component was then transformed to a time constant via the time lag associated with a given coefficient (see Equations 1-2 in **Methods**). In general, this method provides multiple timescales for individual neurons. To assign a single intrinsic timescale to each neuron, we used the longest timescale among all the timescales estimated from statistically significant autoregression coefficients. We did so because dynamics on smaller timescales would reach an asymptote faster and thus are less important for the overall time course of neural response. Intrinsic timescales based on this approach closely match timescales based on autocorrelation (see below).

In addition, because of the structured nature of the task with specific time epochs, we hypothesized that neural response on a given epoch could be influenced by the activity in the same epoch in the previous trials, and thus, we included a second AR component to estimate a “seasonal” timescale for each neuron (**Figure 1a**). The third and fourth types of dependence were captured by two exponential memory-trace components (filters) that could predict fluctuations of each neuron’s response around its average activity profile in a given time bin based on reward feedback and choice on the previous trials, respectively. The corresponding exponential coefficients were then used to estimate the timescale of a “reward-memory” (**Figure 1b**) and “choice-memory” (**Figure 1c**) for each neuron (Bernacchia et al., 2011). Finally, we also included multiple exogenous terms to capture selectivity to reward outcome and choice in the current trial, and their interactions. We used all possible combinations of the two autoregressive and two memory-trace components as well as the presence or absence of exogenous terms (task-relevant signals) to generate 32 (= 2^5^) models (**Supplementary Table 1**).

Considering the complexity of our models, we first tested whether our fitting method could identify the correct model by fitting data generated with one model using all the 32 models (see *Model recovery* in **Methods**). We found that our method could identify the correct model most of the time despite the large number of models considered (**Supplementary Figure 1**). Moreover, we also tested how reliably our methods can identify the best model for each neuron by computing the coefficient of determination (R-squared) for best models and comparing them with those of the second-best models as well as models that only include exogenous terms and thus no timescales.

We found that the best model for each neuron, which often involved about three types of timescales (**Supplementary Figure 2**), captured larger variances of neural activity than the second-best model and the model that did not include any timescales (**Supplementary Figure 3**). These results show that dynamics associated with these timescales indeed capture unique variability in neural response beyond what is predicted by task-relevant signals. Interestingly, the best model for most neurons (∼99.5%) in all four cortical areas included an intrinsic AR component (**Supplementary Figure 4**), illustrating the importance of intrinsic fluctuations in explaining neural variability across cortex. Together, these results demonstrate the robustness of our method in estimating multiple timescales related to dynamics of neural response.

### Parallel but independent hierarchies of timescales

After validating our estimation method and fitting procedure, we used all 32 models to fit individual neurons’ response to identify the best model for each neuron based on cross-validation, and to simultaneously estimate selectivity to task-relevant signals as well as intrinsic, seasonal, choice-memory, and reward-memory timescales. We observed hierarchies for all of the estimated timescales across the four cortical areas, from the lateral intraparietal area (LIP) to the dorsomedial prefrontal cortex (dmPFC) to the dorsolateral prefrontal cortex (dlPFC) to the anterior cingulate cortex (ACC).

The median value of intrinsic timescales increased from ∼60 ms in LIP to ∼120 ms in ACC with the dmPFC and dlPFC exhibiting intermediate values (**Figure 1d**). These intrinsic timescales, however, were significantly smaller than those reported in Murray et al. (2014), which could be due to using the decay on autocorrelations between spikes during the fixation period in that study. To test this possibility, we applied our method to neural response during the fixation period only and found the median intrinsic timescales to be significantly larger for the activity during this epoch compared with the entire trial (**Supplementary Figure 5a**). Nonetheless, applying our method to neural response during the fixation period we observed a range of intrinsic timescales similar to those reported based on autocorrelation.

Similar to intrinsic timescales, our new seasonal timescales also increased from LIP to ACC (**Figure 1e**). However, seasonal timescales were an order of magnitude larger than intrinsic timescales and significantly smaller fractions of neurons exhibited these timescales. Similarly, reward- and choice-memory timescales increased from parietal to prefrontal to cingulate cortex; these timescales assumed values between intrinsic and seasonal timescales (**Figure 1f,g**). Overall, we found that LIP and ACC consistently exhibited the shortest and longest timescales, whereas the two prefrontal areas showed intermediate values. Therefore, our method extended previous findings about intrinsic and reward-memory timescales to the single-cell level and moreover, revealed two new hierarchies of seasonal and choice-memory timescales.

Our results suggest that the estimated timescales increase in tandem across the four cortical areas as is evident from changes in the medians of these timescales across the four areas. To examine this relationship more closely, we computed the correlations between timescales within individual neurons across all cortical areas (cortex-wise correlations) based on simultaneously estimated timescales for each neuron. We found significant correlations between most pairs of timescales except between seasonal and reward-memory timescales and between seasonal and choice-memory timescales (**Figure 2)**. Similar correlation between intrinsic and reward-memory timescales has been reported before but using only population-level estimates (Murray et al., 2014).

**Figure 2.**
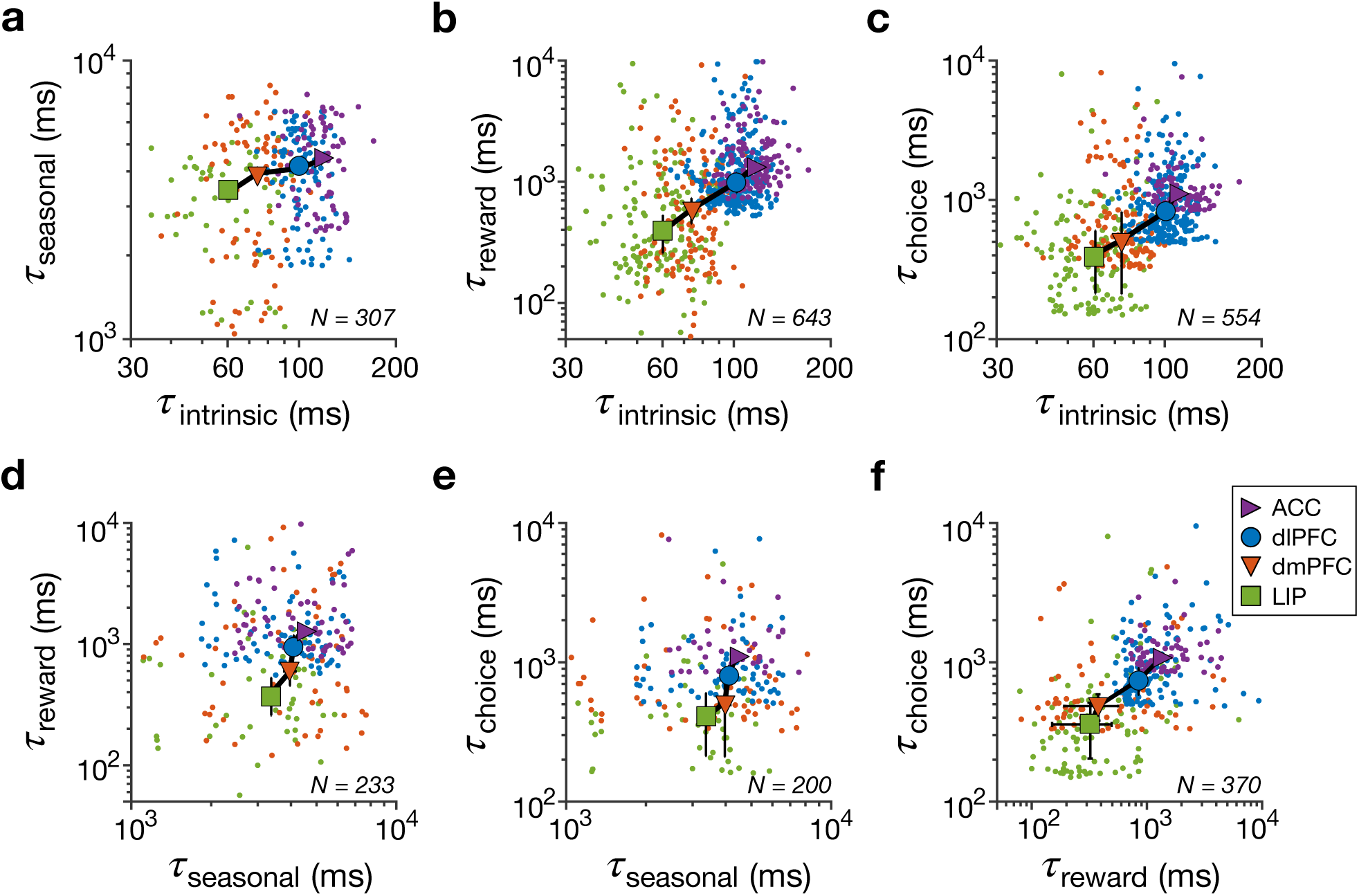
Relationship between different types of timescales across all cortical areas (cortex-wise correlation). Plots show estimated timescales for individual neurons (color dots) and median timescales (symbols) against one another across four cortical areas as indicated in the legend: seasonal vs. intrinsic timescales (**a**), reward-memory vs. intrinsic timescales (**b**), choice-memory vs. intrinsic timescales (**c**), reward-memory vs. seasonal timescales (**d**), choice-memory vs. seasonal timescales (**e**), and choice-memory vs. reward-memory timescales (**f**).Error bars indicate s.e.m. The cortex-wise correlation was significant between most pairs of timescales: intrinsic and seasonal (Spearman correlation, *r* = 0.18, *p* = 0.0013), intrinsic and reward-memory (Spearman correlation, *r* = 0.46, *p* = 0), intrinsic and choice-memory (Spearman correlation, *r* = 0.45, *p* = 6.19 × 10^−14^), and reward-memory and choice-memory (Spearman correlation, *r* = 0.51, *p* = 1.89 × 10^−29^). There was no significant correlation between seasonal and reward-memory timescales (Spearman correlation, *r* = 0.06, *p* = 0.37) and between seasonal and choice-memory timescales (Spearman correlation, *r* = 0.11, *p* = 0.13).

The cortex-wise correlation between timescales could be driven simply by the gradual increase in all timescales across the four cortical areas. Therefore, we tested whether these timescales are correlated across neurons within a given brain area because the presence or lack of correlation between timescales within individual neurons would suggest similar or separate mechanisms for generations of these timescales, respectively. Indeed, we did not find any evidence for correlation between any pairs of timescales in any cortical areas (**Figure 3**). The only evidence for such correlation, which did not survive the Bonferroni correction, was observed between choice- and reward-memory timescales in LIP and dlPFC. Overall, we found that although all four types of timescales consistently increased across cortex in tandem, there was no relationship between them across individual neurons in a given area. This indicates that the previously reported correlation between intrinsic and reward-memory timescales was mostly driven by between-region differences.

**Figure 3.**
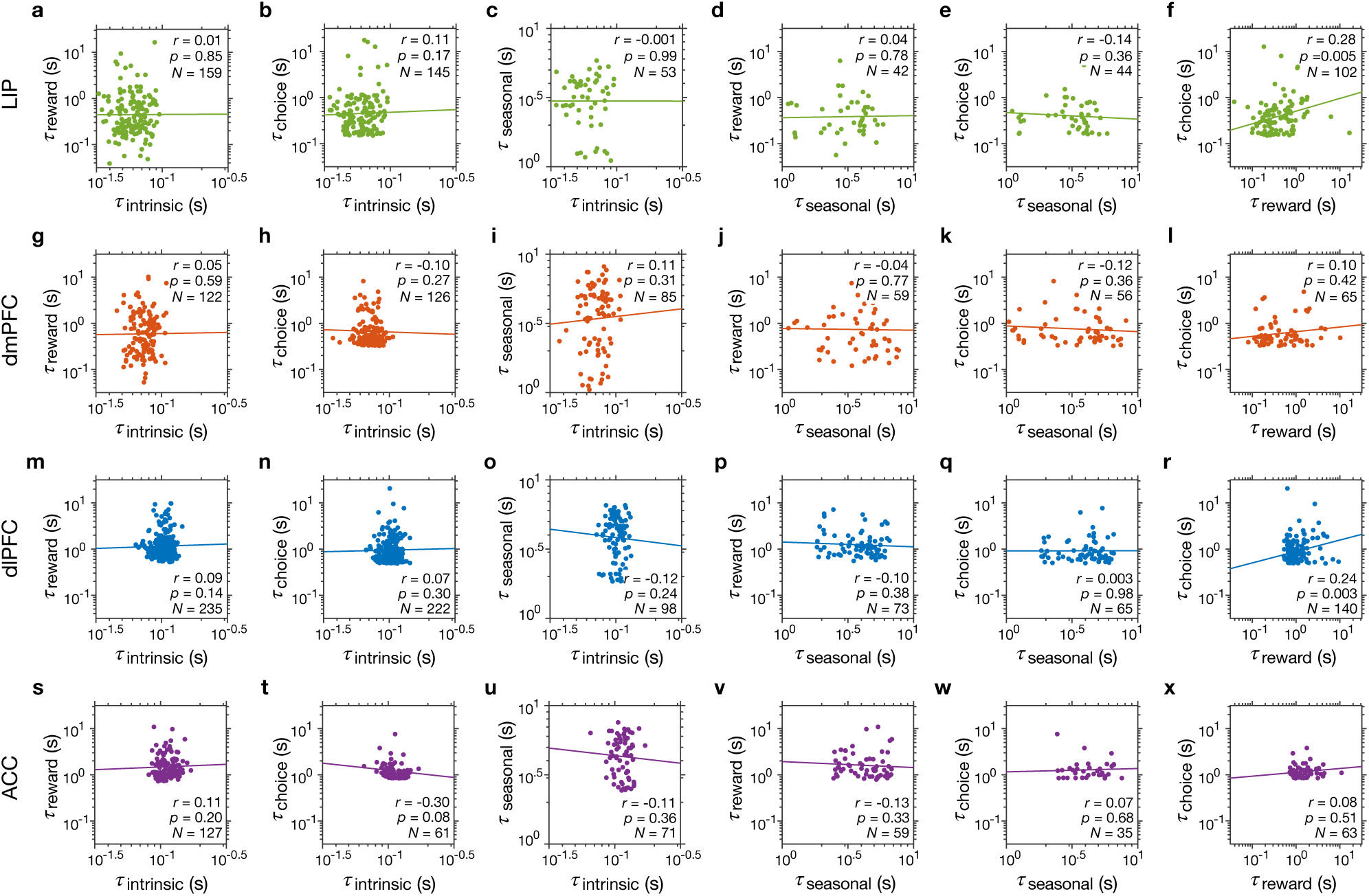
Independence between different types of timescales within individual neurons within individual cortical areas. Each row of panels shows the estimated timescales within individual neurons against one another for a given cortical area indicated on the left. Reported are the Spearman correlation coefficients and corresponding *p*-values, and the number of neurons with significant values of a given pair of timescales. The solid lines represent the regression line that was fit to log timescales.

Considering that these findings are null results, we performed additional simulations to test whether our method is sensitive enough to detect correlations between timescales across individual neurons within a given cortical area if such correlations exist indeed. More specifically, we used activity profiles of randomly selected neurons in our dataset to simulate neural response with significant correlations between certain pairs of timescales, and then used our method to estimate those timescales from the simulated data (see *Correlation recovery simulations* in the **Methods**). We found that our method can detect existing correlations between pairs of timescales and there was no systematic bias in estimated correlations (**Supplementary Figure 6**).

### Behavioral relevance of estimated timescales

To estimate the four timescales, we fit neural response considering all task-relevant signals. This method guarantees that the estimated timescales capture unique variability in neural response, but it is still unclear whether they are relevant and contribute to behavior. To examine whether any of the four neural timescales are relevant for choice behavior during the game of matching pennies, we estimated timescales at which monkeys’ choice behavior on the current trial is influenced by reward and choice on the preceding trials for each session of the experiment (see *Behavioral timescales* in **Methods**). We then calculated the correlations between these two behavioral timescales and each of the four neural timescales.

We found significant correlations between the behavioral reward timescales (which is directly related to the learning rate in the RL models) and reward-memory timescales in all cortical areas (**Figure 4a-d**). A similar relationship has been reported previously but by considering behavioral and neural timescales from three cortical areas together, but has not been tested for individual brain areas (Bernacchia et al., 2011). We also found significant correlations between behavioral choice timescales and choice-memory timescales of neurons in all cortical areas (**Figure 4e-h**). Importantly, there was no significant correlation between the behavioral reward timescales and choice-memory timescales (Spearman correlation; LIP: *r* = 0.16, *p* = 0.052; dmPFC: *r* = −0.05, *p* = 0.60; dlPFC: *r* = −0.06, *p* = 0.37; ACC: *r* = −0.03, *p* = 0.83) or between behavioral choice timescales and reward-memory timescales (Spearman correlation; LIP: *r* = −0.04, *p* = 0.63; dmPFC: *r* = −0.05, *p* = 0.63; dlPFC: *r* = −0.09, *p* = 0.21; ACC: *r* = 0.09, *p* = 0.34). These results illustrate that reward- and choice-memory timescales in all cortical areas were specifically predictive of behavior in terms of the integration of previous choice and reward outcomes, respectively.

**Figure 4.**
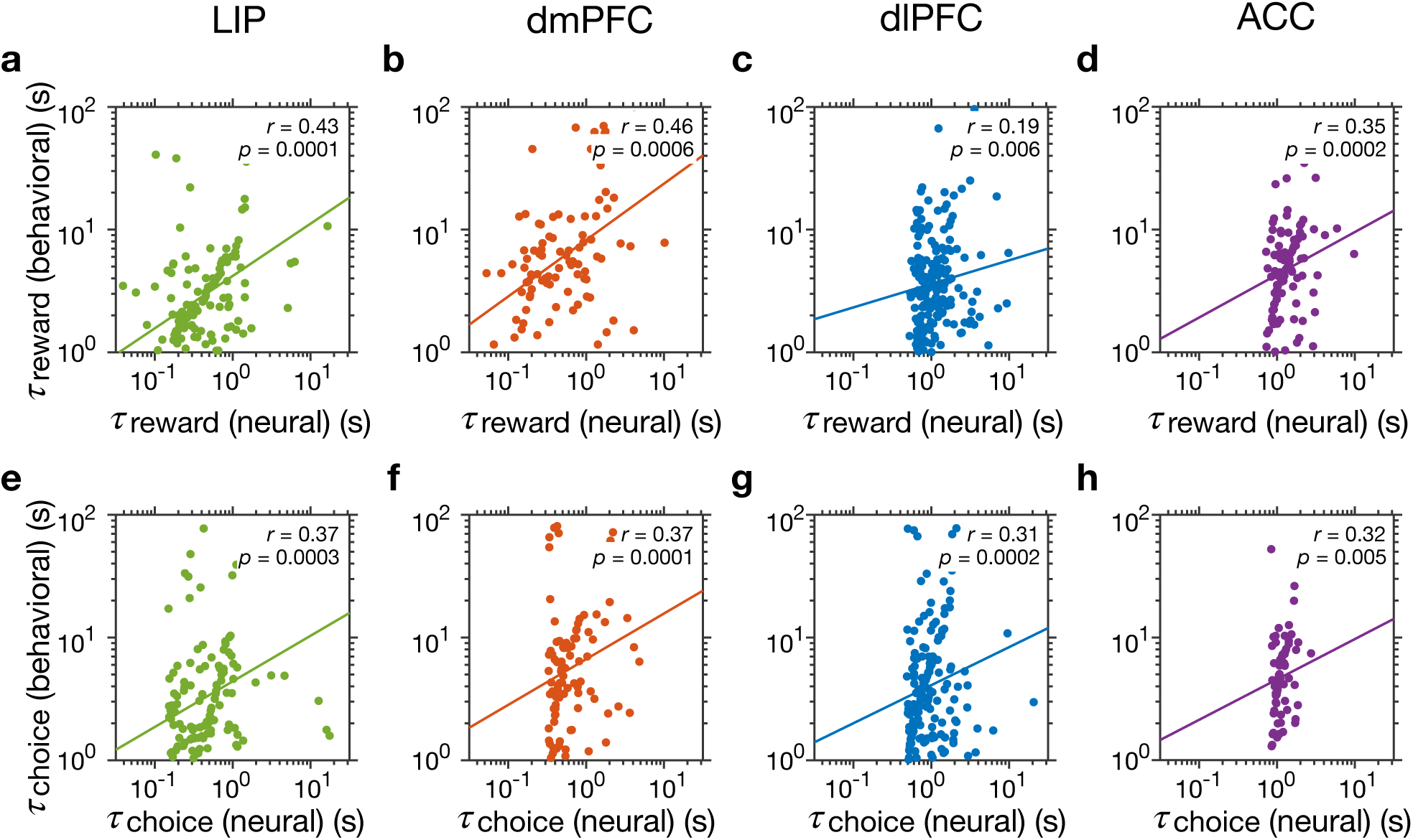
Relationship between reward- and choice-memory timescales and behavioral timescales. (**a–d**) Plots show behavioral reward timescales vs reward-memory timescales of individual neurons recorded during the same sessions, separately for different cortical areas as indicated on the top. Reported are the Spearman correlation coefficients and corresponding *p*-values and the solid lines represent the regression line that was fit to log values. (**e–h**) The same as in a–d but for behavioral choice timescales. There were significant correlations between behavioral and neural timescales in all cortical areas 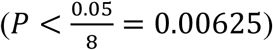.

In contrast to these links between behavioral timescales measuring the decays in the effect of previous reward and choice and corresponding neural timescales, there was no correlation between the behavioral timescales and intrinsic or seasonal timescales (**Supplementary Figure 7**). These results demonstrate that not only our estimated reward-and choice-memory timescales are linked to the integration of reward and choice outcomes over time (trials), but more importantly, intrinsic timescales may not directly contribute to behavior as previously hypothesized (Bernacchia et al., 2011; Murray et al., 2014).

### Dependence of neural timescales on response selectivity

Our finding that four estimated timescales are independent of each other suggests that multiple mechanisms underlie the generation of these timescales. However, if all or some of the estimated timescales depend on response selectivity of individual neurons, inherent heterogeneity in response selectivity could render these timescales decorrelated across individual neurons even if they were generated via a single mechanism. Therefore, we performed additional analyses to examine whether the observed hierarchies of timescales and their relationships depend on the selectivity to task-relevant signals (reward outcome, choice, and their interaction). This was possible because in addition to ensuring that estimated timescales actually captured unique variability in neural response, our method also allowed us to measure the selectivity of individual neurons to task-relevant signals.

First, we found that a significant fraction of neurons in all cortical areas were selective to task-relevant signals, as reflected in the majority of best models to include the exogenous terms (LIP: 66.34%, *χ*^2^ = 11.63, *p* = 9.19 × 10^−5^; dmPFC: 69.73%, *χ*^2^ = 17.56, *p* = 4.73 × 10^−5^; dlPFC: 59.32%, *χ*^2^ = 6.67, *p* = 2.89 × 10^−3^; ACC: 59.74%, *χ*^2^ = 7.72, *p* = 5.17 × 10^−3^). Similar to previous findings based on fits of neural response with regression models (Barraclough et al., 2004; Donahue et al., 2013), we found significant fractions of neurons in all cortical areas to show selectivity to reward outcome right after reward feedback and selectivity to choice after target onset and when a choice was made (**Supplementary Figure 8**). Interestingly, similar fractions of neurons encoded reward outcome across the four cortical areas (*χ*^2^(3) = 3.49, *p* = 0.062) whereas the fraction of neurons selective to choice decreased from LIP to ACC (*χ*^2^(3) = 23.6, *p* = 1.2 × 10^−6^).

Second, we examined whether any of the estimated timescales varied with general or specific selectivity to any task-relevant signals. We did not find any significant difference between timescales of neurons with and without selectivity to task-relevant signals (**Figure 5**; **Supplementary Table 2**). We further examined whether estimated timescales depend on specific selectivity to reward outcome but did not find any evidence for this in any cortical area (**Supplementary Figure 9**; **Supplementary Table 3**). Similarly, we did not find evidence for the dependence of timescales on specific selectivity to choice in any cortical areas; Nonetheless, the overall reward-memory timescales were significantly larger for neurons that were not selective to choice signal (**Supplementary Figure 10**; **Supplementary Table 4**). This suggests a small tendency for neurons with no choice selectivity to integrate reward feedback over longer timescales.

**Figure 5.**
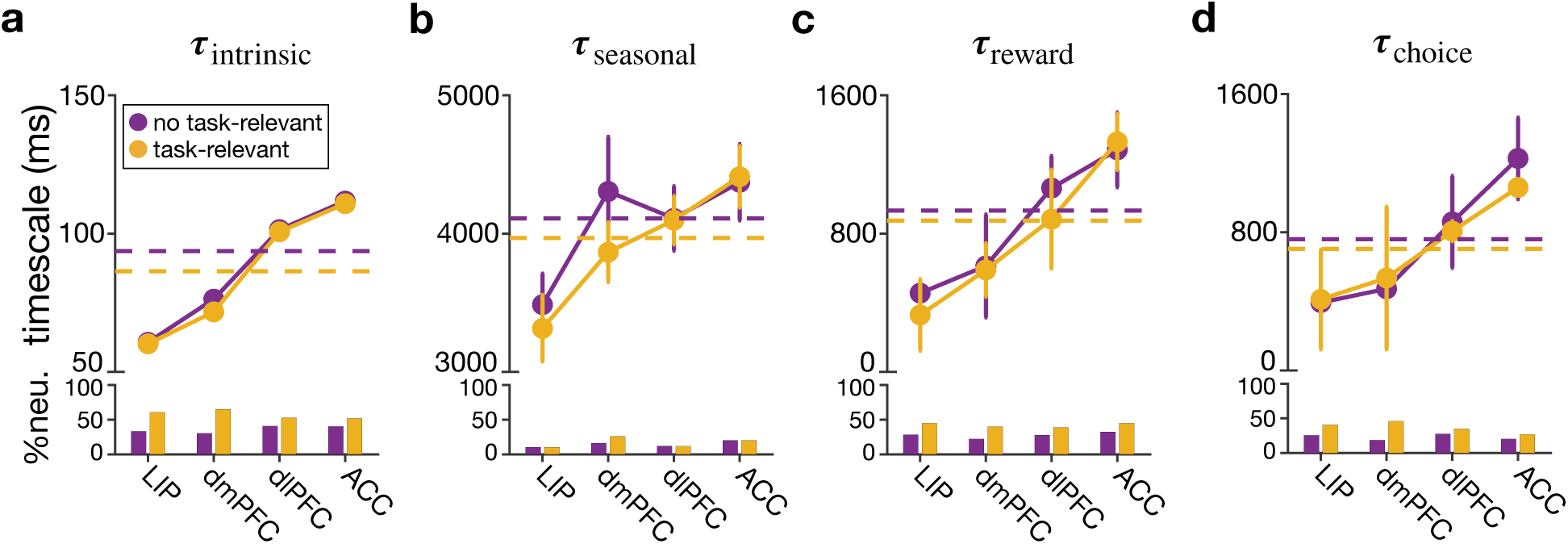
No evidence for the dependence of timescales on overall selectivity to task-relevant (reward outcome and choice) signals. Plots show the median of the estimated intrinsic (**a**), seasonal (**b**), reward-memory (**c**), and choice-memory timescales (**d**) in four cortical areas, separately for neurons with (gold) and without (purple) any type of selectivity to task-relevant signals. The dashed lines show the median across all four areas, and error bars indicate s.e.m. Insets show the fractions of neurons with and without any task-relevant signals (for all neurons with a significant timescale) in different areas. Detailed statistics for comparing neurons with and without task-relevant selectivity are provided in **Supplementary Table 2**.

As mentioned earlier, a few recent studies have shown that the timescales for the decay of autocorrelation in the firing response of individual neurons during the fixation period, often referred to as intrinsic timescales, are predictive of encoding of task-relevant signals. Therefore, we tested whether intrinsic timescales based on autocorrelation depend on selectivity to task-relevant signals. Similar to our results based on our comprehensive method, however, we did not find any difference between the intrinsic timescales of neuron with and without task-relevant selectivity (**Supplementary Figure 5b**). This result indicates that the lack of a relationship between response selectivity and intrinsic timescales is not unique to our method and could be related to the task studied here. Nonetheless, it is important to ensure that intrinsic timescales capture unique variability in neural response; otherwise, any relationship between such timescales and task-relevant signals could be spurious.

Third, despite similar hierarchies of timescales for neurons with different types of selectivity, reward- and choice-memory timescales might still depend on the strength of modulation within each type. To test this possibility, we examined correlations between timescales of reward- and choice-memory integration and the magnitude of selectivity to reward, choice, and their interactions (quantified by standardized regression coefficients for the corresponding exogenous signals) but did not find any significant relationship (**Supplementary Figure 11**). Furthermore, we did not find significant correlation between timescales and neural firing rates in any cortical area (**Supplementary Figure 12**). These results illustrate that activity related to reward and choice memory were independent of immediate response to these signals within individual neurons.

Together, results presented above illustrate the independence of estimated timescales and selectivity to task-relevant signals. These findings suggest that the four estimated timescales related to various dynamics of cortical neural response are not generated via a single mechanism that is then modulated and sculpted by response properties of individual neurons. Instead, they suggest that multiple mechanisms underlie the generation of the four estimated timescales.

## Discussion

We developed a comprehensive and robust method to estimate multiple timescales related to dynamics of neural response along with selectivity of individual neurons to important task-relevant signals. By applying this method to neuronal activity recorded from four cortical areas, we provide evidence for the presence of four parallel timescales related to neural response modulations by previous reward and choice outcomes (reward- and choice-memory timescales), ongoing fluctuations in neural firing (intrinsic timescale), and response during similar task epochs in the preceding trials (seasonal timescale). Although evidence for intrinsic, reward- and choice-memory timescales have been provided before (Bernacchia et al., 2001; Murray et al., 2014), the relationship between these timescales within individual neurons and their dependence on response selectivity and the precise nature of the observed relationships between the selectivity to task-relevant signals and intrinsic timescales based on autocorrelation were not known.

Our method also identified a new timescale related to fluctuations of neural response to experimental epochs (events) across trials and the importance of these dynamics for capturing response variability even though only less than half of the recorded neurons exhibited seasonal timescales. The seasonal timescales were the longest timescales and could reflect internal neural dynamics controlled by top-down signals that could set the state of cortical dynamics that ultimately influence task performance (Carnevale et al., 2012). It is possible that seasonal timescales emerge the task being learned and that is why less neurons exhibit seasonal timescales.

We found four parallel hierarchies of timescales, from parietal to prefrontal to cingulate cortex, at the level of individual neurons. However, none of the four timescales depended on the selectivity of individual neurons to task-relevant signals, and there was no systematic relationship between these timescales across individual neurons in a given cortical area. These indicate that the previously reported correlation between intrinsic and reward-memory timescales (Murray et al., 2014) was mostly driven by between-region differences and was not a property of individual neural response. In addition, our findings contradict a few recent studies showing that intrinsic timescales based on autocorrelation can predict encoding of task-relevant signals in some cortical areas (Cavanagh et al,, 2016; Cavanagh et al., 2018; Fascianelli et al., 2019; Nishida et al., 2014; Wasmuht et al., 2018). This could be due to differences in the experimental paradigms, methods for estimations of intrinsic timescales, or both, and could simply reflect reporting bias considering the large number of studies that have examined the decay in autocorrelation of neural response. More specifically, because intrinsic timescales using the decay in autocorrelation are usually obtained from a fixation period (presumably before strong task-relevant signals emerge in the cortical activity), it is unclear that corresponding dynamics capture unique variability in neural response beyond task-relevant signals, suggesting that the relationship between such intrinsic timescales and encoding of task-relevant signals observed in previous studies might be spurious.

Our results suggest that the four estimated timescales and corresponding dynamics are not generated via a single mechanism and modulated by response properties of individual neurons and, instead, are produced by distinct mechanisms. More specifically, the independence of intrinsic timescales from neural selectivity and the gradual increase of these timescales across cortex confirm the previously postulated role of slower synaptic dynamics (perhaps due to short-term synaptic plasticity) in higher cortical areas (Murray et al., 2014; Wang et al., 2008). Nonetheless, distinct hierarchies of timescales can be generated by other mechanisms than those underlying intrinsic timescales (Miller et al., 2016; Hunt and Hayden, 2017). For example, seasonal timescales could be generated through circuit reverberations evoked by important task events and top-down signals and thus could depend on the dynamics of interactions between neurons in the circuit. Therefore, independence of intrinsic and seasonal timescales within individual neurons challenges the idea that their cortical hierarchies occur due to successive processing of information (Hunt and Hayden, 2017; Yoo and Hayden, 2018) and points to the importance of the heterogeneity of local circuits to which a neuron belongs (Chaudhuri et al., 2015). That is, a neuron could receive synaptic inputs with fast dynamics (e.g., due to short-term plasticity) but contributes to slow circuit reverberations that generate seasonal timescales, and vice versa.

We found that choice- and reward-memory timescales selectively predict behavioral timescales related to behavioral integration of previous choice and reward outcomes, respectively and thus, are relevant to choice behavior. This indicates that these timescales are more likely to depend on long-term reward- and choice-dependent synaptic plasticity as presumed in different reinforcement learning models. Assuming Hebbian form of synaptic plasticity, one could predict that stronger response to reward feedback should result in stronger changes in synaptic plasticity and thus a shorter timescale for reward memory for a given neuron. However, we did not find any evidence for a relationship between these memory timescales and response selectivity to reward and choice outcomes on the current trial. This result could indicate the presence of significant heterogeneity in synaptic plasticity rules. In addition, reward and choice-memory timescales could be decorrelated because choice and reward are only weakly coupled during the game of matching pennies, but this might also reflect the fact that synaptic plasticity depends on activity and reward history (Abraham, 2008; Farashahi et al., 2017). More specifically, because synaptic plasticity changes with preceding neural activity and reward history, heterogeneity in both of these factors could make choice- and reward-memory decorrelated. Finally, the lack of correlation between intrinsic (and seasonal) timescales and reward- or choice-memory timescales could undermine the proposal that intrinsic dynamics directly contribute to reward-based and goal-directed behavior (Murray et al., 2014). Instead, we speculate that the independence of different timescales within individual neurons could contribute to behavioral flexibility by allowing neurons to integrate different pieces of task-relevant information independently (Farashahi et al., 2019).

Although we only investigated response dynamics of individual neurons, there are recent studies showing that population activity exhibits similar dynamics and reflect task-relevant information (Kobak et al., 2016; Cueva et al., 2019; Rossi-Pool et al., 2019). For example, Kobak and colleagues showed that after proper de-mixing and dimensionality reduction, population response reflects task parameters such as reward and choice, similar to response of single neurons (Kobak et al., 2016). In another study, Rossi-Pool and colleagues found that during a temporal pattern discrimination task, population activity in the dorsal premotor cortex exhibit temporal dynamics similar to those of single neurons, but these dynamics diminish during a non-demanding task (Rossi-Pool et al., 2019). Future studies are needed to compare the timescales underlying response of individual neurons to those of populations of neurons, which could be important for understanding how corresponding dynamics are generated.

## Methods

### Neural data

All experimental procedures followed the guidelines by the US National Institutes of Health and were approved by the University Committee on Animal Research (UCAR) at the University of Rochester and the Institutional Animal Care and Use Committee (IACUC) at Yale University. Experimental details for the data sets have been reported previously (Barraclough et al., 2004; Seo et al., 2009; Seo and Lee, 2007; Donahue et al., 2013). We used single-neuron spike train data that were recorded in macaque monkeys performing a competitive decision-making task of matching pennies. In each trial, monkeys chose one of two color targets by shifting their gaze while the computer made its choice by simulating a rational opponent; the animal received reward if its choice matched that of the computer (Barraclough et al., 2004). We used spike counts in 50-msec time bins starting with reward feedback (post-feedback) and spanning into the following trial (there could be a maximum of 80 bins in a given trial). This choice of starting point was only for computational convenience. Data includes recordings from 205 neurons in the lateral intraparietal area (LIP) from 1 female and 2 male monkeys (Seo et al., 2009), 185 neurons in the dorsomedial prefrontal cortex (dmPFC) from 2 male monkeys (Donahue et al., 2013), 322 neurons in the dorsolateral prefrontal cortex (dlPFC) from 1 female and 4 male monkeys (Barraclough et al., 2004), and 154 neurons in the dorsal anterior cingulate cortex (ACC) from 2 male monkeys (Seo and Lee, 2007).

### Model

Our goal here was to predict the spike counts as a non-stationary time series based on the preceding neural activity and task-relevant signals in order to simultaneously estimate various timescales in neural response and selectivity to task-relevant signals. A powerful method to capture non-stationarity of a time series due to factors with different timescales is the seasonal autoregressive with exogenous inputs (ARX) model, which commonly has been used in various fields (Seo and Lee, 2007; Seo et al., 2007, Hipel and McLeod, 1994; Hamzaçebi, 2008; Box et al., 2015). The autoregressive component of the ARX model aims to predict the output variable or response based on immediately preceding response, whereas the seasonal component allows the model to capture fluctuations or variations due to the periodic nature of the external factors. In the context of neural data of our experiments, the seasonal component refers to the relationship between neural response across trials due to the specific structure of the task (see below).

To predict neural response, we included two autoregressive components in our model, resulting in a seasonal 2D-ARX model that also includes two exponential memory traces for choice and reward. First, we assumed that neural response at any time point in a trial depends on earlier activity in the same trial (Murray et al., 2014). This dependence was captured by an autoregressive component that predicts spikes in a given 50-msec time bin based on spikes in the preceding *F* time bins (autoregressive model with order *F*). Therefore, this “intrinsic” autoregressive component (referred to as AR_intrinsic_) uses a weighted average of firing rates in the preceding bins in order to predict the current firing rate. Our preliminary results showed that there is more than one significant autoregression coefficient for most neurons. In order to assign a single *intrinsic* timescale (*τ*_*intrinsic*_) for each neuron, we selected the longest timescale among the AR_intrinsic_ coefficients (see Equations 1-3 below) because neural dynamics on smaller timescales would reach an asymptote and thus, are less important for the overall time course of neural response. We found that in addition to the longest timescales closely matching timescales based on autocorrelation (**Supplementary Figure 5**), the second longest timescales also exhibit a similar hierarchy but on a smaller range (data not shown).

Second, because of the structured nature of the task with specific time epochs, we hypothesized that neural response on a given epoch could be influenced by the activity in the same epoch in the preceding trials. In other words, task epochs could provide a “seasonal” source of variability in neural response. Therefore, we included a seasonal autoregressive component (referred to as AR_seasonal_) in our model in order to predict response in the current time bin based on responses in the same time bins in the preceding *G* trials (autoregressive model with order *G*; Equation 1). The corresponding autoregression coefficients were used to estimate a *seasonal* memory timescale (*τ*_*seasonal*_) for each neuron. Therefore, seasonal timescales capture how fluctuations of activity during a given epoch decay over trials (more precisely, time difference between the same epoch over successive trials).

Third, we assumed that neural response at any time point in a trial depends on reward outcome (reward vs. no reward) and choice (left vs. right choice) in the preceding trials (Bernacchia et al., 2011). These dependences relate spikes in a given time bin of the current trial to reward and choice outcomes in the preceding H trials. To capture these dependencies, we assumed two separate exponential memory-trace filters that are modulated by the average response in a given epoch and previous reward or choice signals (see Equation 1 below). The corresponding exponential memory-trace coefficients were used to estimate a *reward-memory* timescale (*τ*_*reward*_) and a *choice-memory* timescale (*τ*_*choice*_) for each neuron. Therefore, reward- and choice-memory timescales capture how the influence of reward and choice outcomes decays over time (time from the preceding reward feedback and choice, respectively).

Finally, we also included various exogenous terms to capture selectivity in response to current choice (*C*), current reward (*R*), and their interaction (*C* × *R*). We did not include terms for previous choice and reward because effects of previous choice and reward are captured by choice- and reward-memory, respectively. The selectivity to task-relevant signals was captured using four boxcars relative to relevant events in the task. More specifically, we considered: a) three terms (regressors) for choice, one for [0,500] msec interval relative to the onset of choice targets (Choice 1), one for [0,500] msec interval relative to target fixation (Choice 2), and one for [0,500] msec interval relative to reward feedback (Choice 3); b) one term for reward for [0,500] msec interval relative to reward feedback; and c) one term for the interaction of choice and reward for [0,500] msec interval relative to reward feedback.

Based on the description provided above, the model involves weighted average over 2D space of preceding time bins and trial firing rates (with order *F* for time bins and *G* for trials) in addition to two separate weighted averages over reward and choice outcomes on the preceding *H* trials. More formally, the spike counts in bin *n* of trial *k, y*(*n, k*) is given by the following equation:

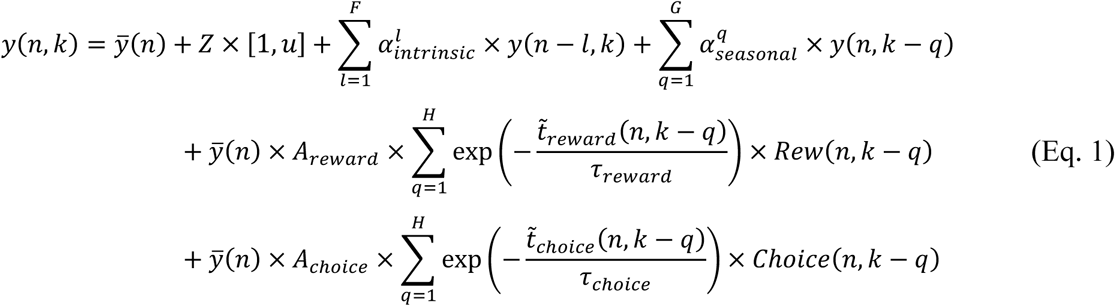

where 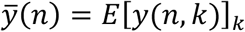 is the average value of spike count in each bin over all trials, *Z*(1 × 6) is the vector of coefficients for the task-relevant component, and *u* is a row vector of 5 task-relevant inputs (three choice signals, reward, and interaction of choice and reward).

Autoregressive coefficients for intrinsic and seasonal fluctuations are denoted by 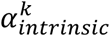 and 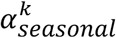, respectively 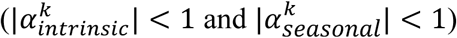. Reward- and choice-memory timescales are indicated by *τ*_*reward*_ and *τ*_*choice*_ whereas *A*_*reward*_ and *A*_*choice*_ are the amplitudes of reward- and choice-memory components, respectively. 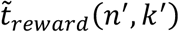 and 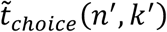 indicate the time difference between the time bin (with time resolution of 50 msec) at which reward or choice was occurred right before time bin (*n*′) on trial (*k*′) and current time. Finally, we used AR_intrinsic_ and AR_seasonal_ of order 5 (*F* = *G* = 5) and computed reward- and choice-memory traces over the preceding 5 trials (*H* = 5).

For an autoregressive model of order 1, AR(1), that only has one single autoregressive coefficient (*y*(*t*) = ø_1_*y*(*t* − 1)), a single timescale can be defined equal to 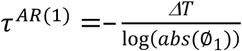, where *ΔT* is the size of the time bin (time resolution). By extending the same logic, we defined a set of intrinsic (*τ*_*intrinsic*_) and seasonal (*τ_seasonal_*) timescales based on the AR components as follows:

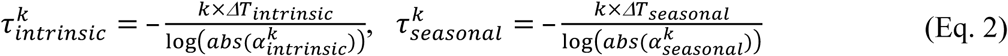

Therefore, the timescales of the AR components depend on both autoregressive components and the length of time lags for bins (*ΔT*_*intrinsic*_ = 50 *msec*) and trials (*ΔT_seasonal_*: average trial length). Because the AR_intrinsic_ and AR_seasonal_ components provide multiple timescales, we used the longest timescale to assign a single intrinsic and seasonal timescales to each neuron:

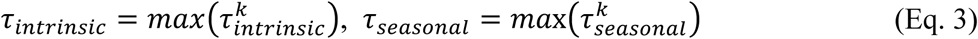

Note that we only considered AR coefficients that were statistically significant for computing the corresponding timescales.

This is the most general model from which we constructed more specific models by turning on and off the autoregressive components, reward- and choice-memory traces, and task-relevant terms. These constructions resulted in 32 alternative models, consisting of all the possible combinations of the general model components (the list of all possible combinations of the models can be found in **Supplementary Table 1**). To select the best model for each neuron, we ran all 32 models and used cross-validation (see Model selection and parameters below).

Two special cases of our model could mimic the autocorrelation model of Murray et al. (2014) and the exponential memory-traces model of Bernacchia et al. (2011) using only the AR_intrinsic_ and reward-memory components, respectively. To replicate the results of the two previous studies, we used their methods for profiling neural activity. More specifically, Murray et al. (2014) used spike counts in a period starting from fixation point to 500 msec after that (post-fixation), and then split spikes in this post-fixation epoch into 10 time bins of 50 msec. Bernacchia et al. (2011) used two time periods, each spanning 1500 msec, that were further divided into 6 time bins of 250 msec each. The first period consisted of 6 successive, 250 msec bins starting from 1000 msec before target onset to 500 msec after that. The second period consisted of 6 successive, 250 msec bins starting from 500 msec before feedback period to 1000 msec after that.

### Model selection and parameters

Model parameters for the autoregressive components were determined by finding the best model for each neuron based on their performance (using R-squared measure). We performed a ten-fold cross-validation fitting process to calculate the overall performance for each model. Specifically, we generated each instance of training data by randomly sampling 90% of all data (bins) for each neuron and then calculated fitting performance based on R-squared in the remaining 10% of data (test data). This process was repeated 30 times, and the performance was computed based on the median of performance across these 30 instances. To identify the best fit for each instance of the training data, we ran the model 50 times from different initial parameter values and minimized the residual sum of squares to obtain the best model parameters. The median of model parameters over the 30 instances was used to compute the best parameters for each model. In order to be able to compare parameters across different cross-validation instances, we z-scored all input and output vectors before fitting each instance.

To remove the outlier model parameters in a given cortical area, we used 1.5×IQR method for each parameter (and not neuron). In order to determine the type of selectivity to task-relevant signals for each neuron, we first identified neurons for which the model with exogenous terms provided the better fit. Neurons with a significant parameter value for a given task-relevant signal (e.g., reward signal) were determined as the neurons with that type of task-relevant selectivity (e.g., reward-selective neurons).

### Model recovery

In order to examine whether our method is able to identify the correct model, we measured the probability of finding the correct model in the simulated data. More specifically, we generated 500 sets of simulated neural data based on a given model and the actual activity profiles of recorded neurons, and then fit those data with all 32 models. We used the goodness-of-fit based on the AIC to determine the best model.

More specifically, to generate alternative profiles of neural activity (neural profiles), we randomly selected 500 average neural response (divided into 50-msec time bins) from the 866 available neurons in the four areas. We used the original neural profiles for 300 out of 500 neurons and generated synthetic profiles from the remaining 200 profiles by permuting blocks of bins (5 bins in each block) of the original neural profiles. We then generated a set of 5000 values for the four types of timescales using the estimated range of timescales across all areas. For each neural profile, we used 10 randomly selected timescale values to produce spike counts in each bin.

### Correlation recovery simulations

To show that our method is sensitive enough to detect correlations between timescales of individual neurons within each cortical area, given such correlations exist, we performed the following simulations. First, we randomly selected 100 activity profiles (i.e., mean neural response from individual neurons) from neurons in our dataset. We then assigned a random set of four timescales to each profile and tested whether a certain pair of timescales (e.g., *τ_intrinsic_* and *τ_seasonal_*) are significantly correlated (with 0.05 ≤ *abs*(*ρ*) ≤ 0.75) across the 100 profiles by chance. If so, we used our full ARMAX model to generate spike counts using the activity profiles and the chosen timescales. We repeated this procedure 60 times in order to generate 60 datasets of neural response for which there is a significant correlation between a given pair of timescales. We then used our full model to estimate timescales for neural response in each generated dataset and subsequently tested correlation between the estimated timescales.

### Estimation of behavioral timescales

In order to estimate behavioral timescales related to the influence of previous choice and reward outcomes we used a simple RL model with two sets of values functions that are updated according to reward outcomes and choice on every trial. More specifically, the reward-dependent value functions for choosing target *x* (Left or Right option) on trial *t, Q*_*t*_(*x*), is updated according to the following equation (Sutton and Barto, 1998) as follows:

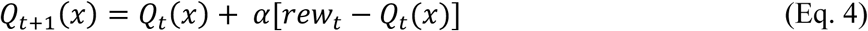

where *rew*_*t*_ (equal to 1 or 0) is the reward received by the animal on trial *t*, and *α* is the learning rate. This update rule can be rearranged as:

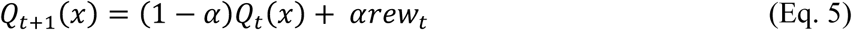

to show that 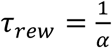 can be used as the behavioral memory timescale of previous reward outcomes. We also considered a set of two choice-dependent value functions for capturing the effect of previous choices over time:

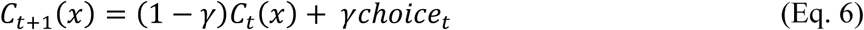

where *C*_*t*_(*x*) denotes the choice-dependent value function for target *x* on trial *t, choice*_*t*_ is the choice on trial *t* (Left or Right) and *γ* is the decay rate. Similar to behavioral reward memory, 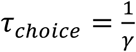 can be used as the behavioral memory timescale of previous choice outcomes.

Importantly, overall value of selecting target *x* on trial *t, V*_*t*_(*x*), is a linear sum of the two value functions:

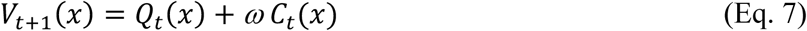

where *ω* determines the relative contribution of choice-dependent component. Finally, the probability that the animal chooses the rightward target on trial *t, P*_*t*_(*R*), was determined using a softmax function of the overall values:

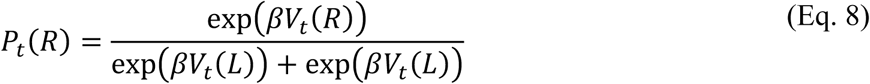

We used this model to fit choice data in each recording session of the experiment to estimate model parameters (*α, γ, ω, β*) using a maximum likelihood procedure.

## Acknowledgments

This work is supported by the National Institutes of Health (grant R01 DA047870 to AS, and grants R01 DA029330 and R01 MH 108629 to DL).

---

**Supplementary Table 1.** List of all possible combinations of the models used to fit neural response and their main components. Exogenous terms refer to regressors included in the model to capture the effects of task-relevant signals on neural response.

| Model # | Model components |
| --- | --- |
| 1 | average response only |
| 2 | exogenous terms only |
| 3 | intrinsic autoregressive only |
| 4 | exogenous terms and intrinsic autoregressive |
| 5 | seasonal autoregressive only |
| 6 | exogenous terms and seasonal autoregressive |
| 7 | intrinsic and seasonal autoregressive |
| 8 | exogenous terms, intrinsic and seasonal autoregressive |
| 9 | reward-memory trace only |
| 10 | exogenous terms and reward-memory trace |
| 11 | intrinsic autoregressive and reward-memory trace |
| 12 | exogenous terms, intrinsic autoregressive and reward-memory trace |
| 13 | seasonal autoregressive and reward-memory trace |
| 14 | exogenous terms, seasonal autoregressive and reward-memory trace |
| 15 | intrinsic and seasonal autoregressive plus reward-memory trace |
| 16 | exogenous terms, intrinsic and seasonal autoregressive plus reward-memory trace |
| 17 | choice-memory trace only |
| 18 | exogenous terms and choice-memory trace |
| 19 | intrinsic autoregressive and choice-memory trace |
| 20 | exogenous terms, intrinsic autoregressive and choice-memory trace |
| 21 | seasonal autoregressive and choice-memory trace |
| 22 | exogenous terms, seasonal autoregressive and choice-memory trace |
| 23 | intrinsic and seasonal autoregressive plus choice-memory trace |
| 24 | exogenous terms, intrinsic and seasonal autoregressive plus choice-memory traces |
| 25 | reward- and choice-memory traces |
| 26 | exogenous terms plus reward- and choice-memory traces |
| 27 | intrinsic autoregressive plus reward- and choice-memory traces |
| 28 | exogenous terms, intrinsic autoregressive plus reward- and choice-memory traces |
| 29 | seasonal autoregressive plus reward- and choice-memory traces |
| 30 | exogenous terms, seasonal autoregressive plus reward- and choice-memory traces |
| 31 | intrinsic and seasonal autoregressive plus reward- and choice-memory traces |
| 32 | exogenous terms, intrinsic and seasonal autoregressive plus reward- and choice-memory traces |

**Supplementary Table 2.** Dependence of timescales on overall selectivity to task-relevant (reward outcome and choice) signals. Reported are *p*-values (two-sided Wilcoxon ranksum test) and effect sizes for the difference in timescales between neurons selective to task-relevant signals and those not selective to task-relevant signals in a given cortical area, and all areas combined. There was no significant difference between estimated timescales of the two types of neurons 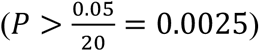.

| **Timescales** | **Cortical area** | ***p*-value** | **Cohen's *d*** | **d.f.** | **#task-relevant** | **#non task-relevant** |
| --- | --- | --- | --- | --- | --- | --- |
| Intrinsic timescales | LIP | 0.88 | 0.01 | 111.29 | 124 | 68 |
|  | dmPFC | 0.07 | 0.14 | 91.67 | 120 | 56 |
|  | dIPFC | 0.55 | 0.03 | 226.84 | 170 | 131 |
|  | ACC | 0.90 | 0.01 | 113.51 | 80 | 62 |
|  | **All** | 0.07 | 0.09 | 654.39 | 494 | 317 |
| Seasonal timescales | LIP | 0.58 | 0.08 | 36.86 | 21 | 21 |
|  | dmPFC | 0.16 | 0.16 | 49.41 | 48 | 30 |
|  | dIPFC | 0.73 | 0.04 | 74.82 | 38 | 39 |
|  | ACC | 0.57 | 0.07 | 59.66 | 31 | 31 |
|  | **All** | 0.28 | 0.07 | 246.26 | 138 | 121 |
| Reward-memory timescales | LIP | 0.43 | 0.06 | 108.32 | 92 | 58 |
|  | dmPFC | 0.97 | 0.003 | 64.32 | 74 | 41 |
|  | dIPFC | 0.02 | 0.15 | 193.79 | 125 | 89 |
|  | ACC | 0.88 | 0.01 | 102.09 | 69 | 50 |
|  | **All** | 0.18 | 0.05 | 595.12 | 360 | 238 |
| Choice-memory timescales | LIP | 0.88 | 0.01 | 115.94 | 83 | 52 |
|  | dmPFC | 0.12 | 0.14 | 98.10 | 85 | 34 |
|  | dIPFC | 0.50 | 0.05 | 97.63 | 112 | 89 |
|  | ACC | 0.06 | 0.22 | 32.92 | 41 | 31 |
|  | **All** | 0.52 | 0.03 | 510.35 | 321 | 206 |

**Supplementary Table 3.**
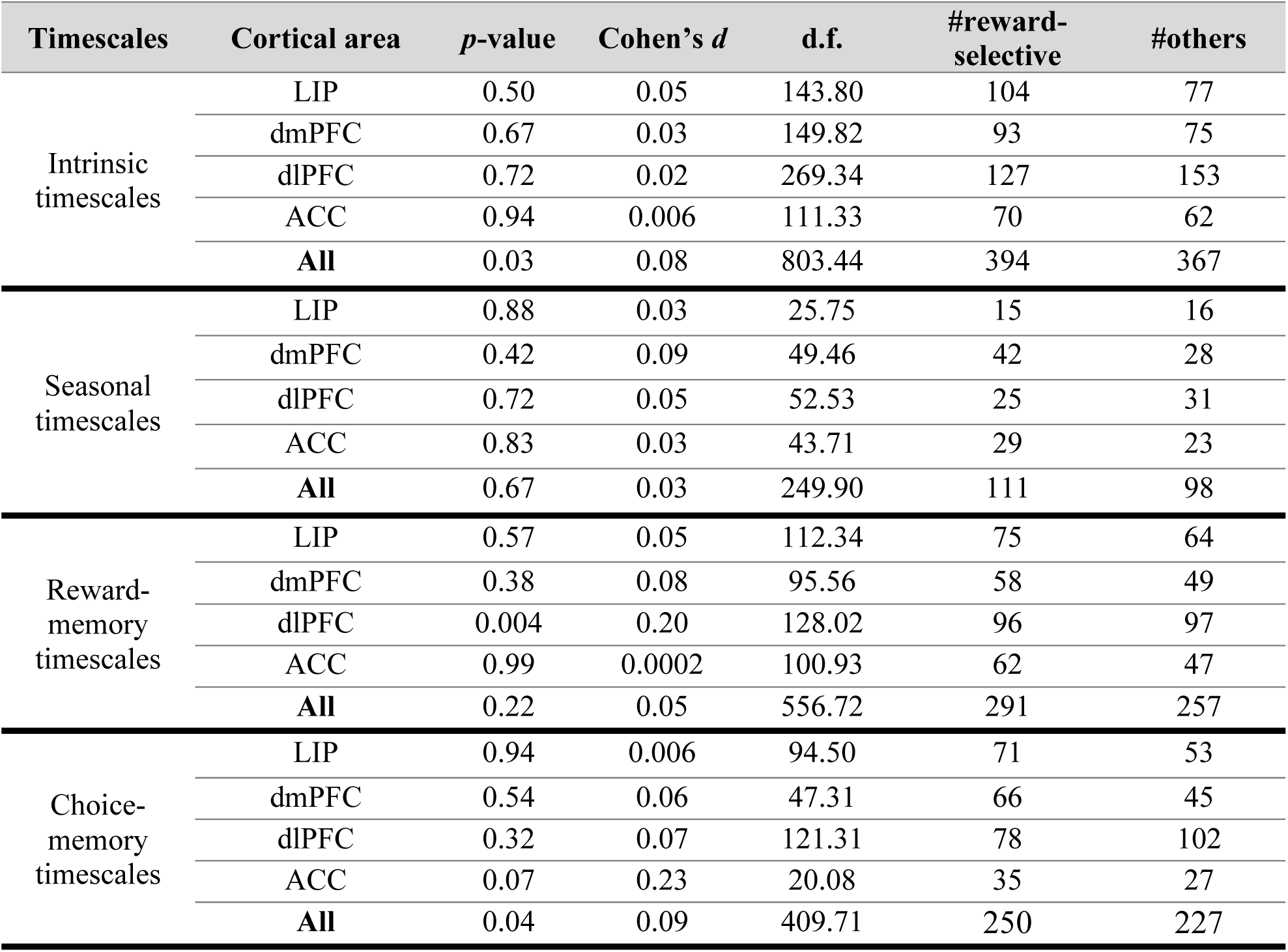
Comparisons of timescales between neurons with selectivity to reward and those with other types of selectivity. Reported are *p*-values (two-sided Wilcoxon ranksum test) and effect sizes for the difference in estimated timescales between reward and non-reward selective (i.e., those selective to choice or interaction of reward and choice) neurons, separately for each cortical area and across all areas. There was no significant difference between neurons selective to reward and the non-reward signals in a given area or across all areas 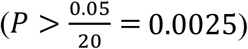.

| Timescales | Cortical area | *p*-value | Cohen's *d* | d.f. | #reward-selective | #others |
| --- | --- | --- | --- | --- | --- | --- |
| Intrinsic timescales | LIP | 0.50 | 0.05 | 143.80 | 104 | 77 |
|  | dmPFC | 0.67 | 0.03 | 149.82 | 93 | 75 |
|  | dIPFC | 0.72 | 0.02 | 269.34 | 127 | 153 |
|  | ACC | 0.94 | 0.006 | 111.33 | 70 | 62 |
|  | **All** | 0.03 | 0.08 | 803.44 | 394 | 367 |
| Seasonal timescales | LIP | 0.88 | 0.03 | 25.75 | 15 | 16 |
|  | dmPFC | 0.42 | 0.09 | 49.46 | 42 | 28 |
|  | dIPFC | 0.72 | 0.05 | 52.53 | 25 | 31 |
|  | ACC | 0.83 | 0.03 | 43.71 | 29 | 23 |
|  | **All** | 0.67 | 0.03 | 249.90 | 111 | 98 |
| Reward-memory timescales | LIP | 0.57 | 0.05 | 112.34 | 75 | 64 |
|  | dmPFC | 0.38 | 0.08 | 95.56 | 58 | 49 |
|  | dIPFC | 0.004 | 0.20 | 128.02 | 96 | 97 |
|  | ACC | 0.99 | 0.0002 | 100.93 | 62 | 47 |
|  | **All** | 0.22 | 0.05 | 556.72 | 291 | 257 |
| Choice-memory timescales | LIP | 0.94 | 0.006 | 94.50 | 71 | 53 |
|  | dmPFC | 0.54 | 0.06 | 47.31 | 66 | 45 |
|  | dIPFC | 0.32 | 0.07 | 121.31 | 78 | 102 |
|  | ACC | 0.07 | 0.23 | 20.08 | 35 | 27 |
|  | **All** | 0.04 | 0.09 | 409.71 | 250 | 227 |

**Supplementary Table 4.** Comparisons of timescales between neurons with selectivity to choice and those with other types of selectivity. Reported are *p*-values (two-sided Wilcoxon ranksum test) and effect sizes for the difference in estimated timescales between choice and non-choice selective (i.e., those selective to reward or interaction of reward and choice) neurons, separately for each cortical area and across all areas. Orange shading indicates a significant effect. There was only a significant difference between reward-memory timescales for neurons selective to choice and the non-choice signals across all areas 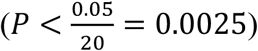.

| Timescales | Cortical area | *p*-value | Cohen's *d* | d.f. | #choice-selective | #others |
| --- | --- | --- | --- | --- | --- | --- |
| Intrinsic timescales | LIP | 0.32 | 0.07 | 148.25 | 106 | 75 |
|  | dmPFC | 0.01 | 0.20 | 119.82 | 105 | 63 |
|  | dIPFC | 0.18 | 0.08 | 266.59 | 146 | 134 |
|  | ACC | 0.17 | 0.12 | 43.89 | 28 | 104 |
|  | **All** | 0.004 | 0.21 | 800.19 | 385 | 376 |
| Seasonal timescales | LIP | 0.53 | 0.11 | 28.81 | 18 | 13 |
|  | dmPFC | 0.19 | 0.16 | 44.50 | 43 | 27 |
|  | dIPFC | 0.87 | 0.02 | 55.31 | 29 | 27 |
|  | ACC | 0.32 | 0.14 | 4.83 | 5 | 47 |
|  | **All** | 0.09 | 0.10 | 254.68 | 95 | 114 |
| Reward-memory timescales | LIP | 0.54 | 0.05 | 104.27 | 81 | 58 |
|  | dmPFC | 0.96 | 0.005 | 80.48 | 64 | 43 |
|  | dIPFC | 0.02 | 0.17 | 160.20 | 106 | 87 |
|  | ACC | 0.69 | 0.04 | 24.37 | 21 | 88 |
|  | **All** | 0.0007 | 0.14 | 464.38 | 272 | 276 |
| Choice-memory timescales | LIP | 0.27 | 0.10 | 83.44 | 68 | 56 |
|  | dmPFC | 0.19 | 0.12 | 38.62 | 73 | 38 |
|  | dIPFC | 0.46 | 0.05 | 92.22 | 98 | 82 |
|  | ACC | 0.93 | 0.01 | 57.25 | 12 | 50 |
|  | **All** | 0.21 | 0.05 | 409.97 | 251 | 226 |

**Supplementary Figure 1.**
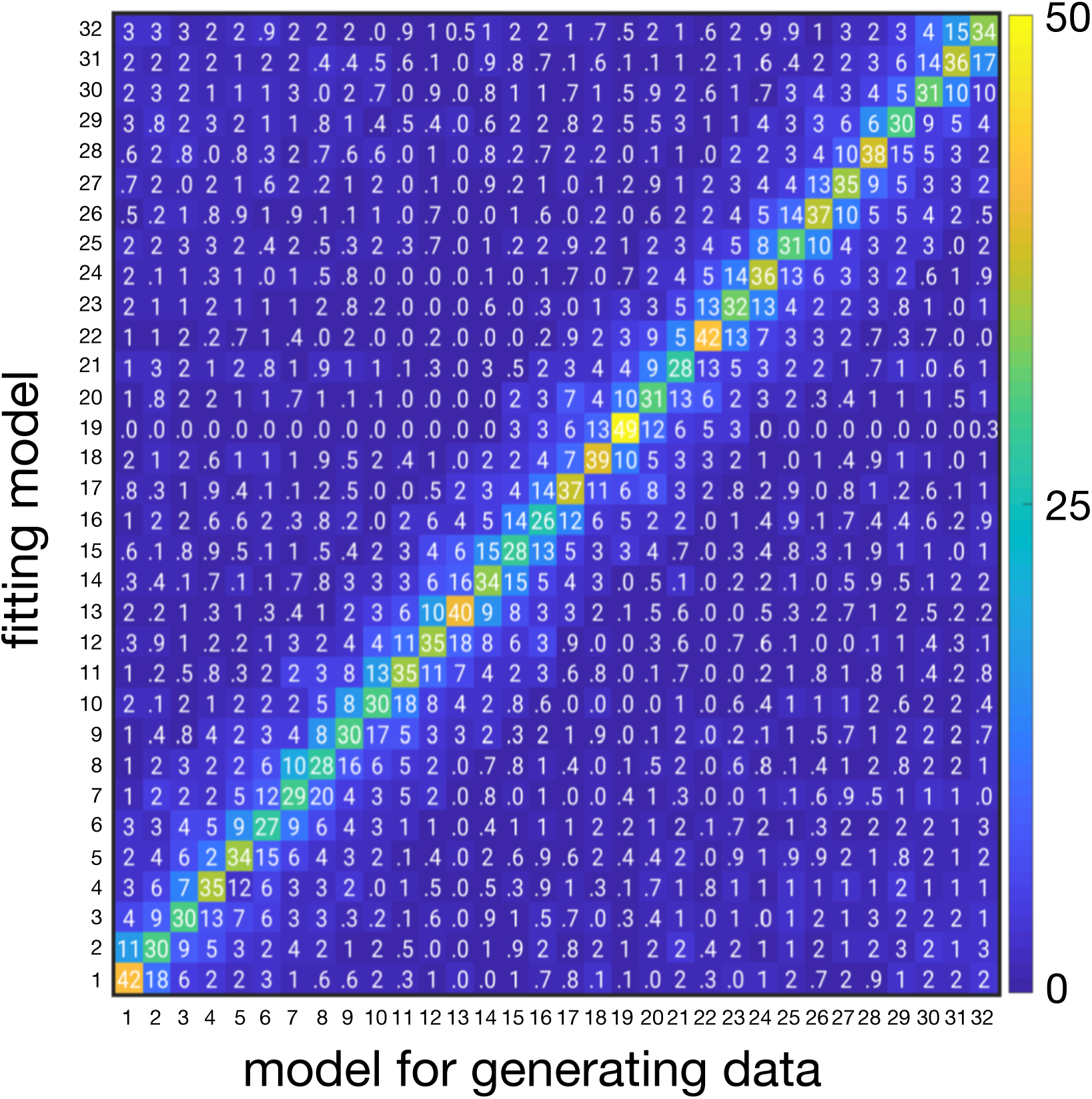
Model recovery. Our method is able to identify the correct model. The value of each cell reports the percentage of instances that a model used to generate the data (shown on the x-axis) was best fit by another model (fitting model, shown on the y-axis). The model corresponding to each number is provided in **Supplementary Table 1**. The model with the minimum AIC was assigned as the best model. The probability for assigning the best model by chance is ∼3% and thus, values above 25% on the diagonal indicate that in most cases the correct model was identified. For these simulations, we generated 500 sets of simulated neural data based on a given model and actual activity profile of recorded neurons, and then fit those data with all the models in order to calculate the AIC and determine the best model.

**Supplementary Figure 2.**
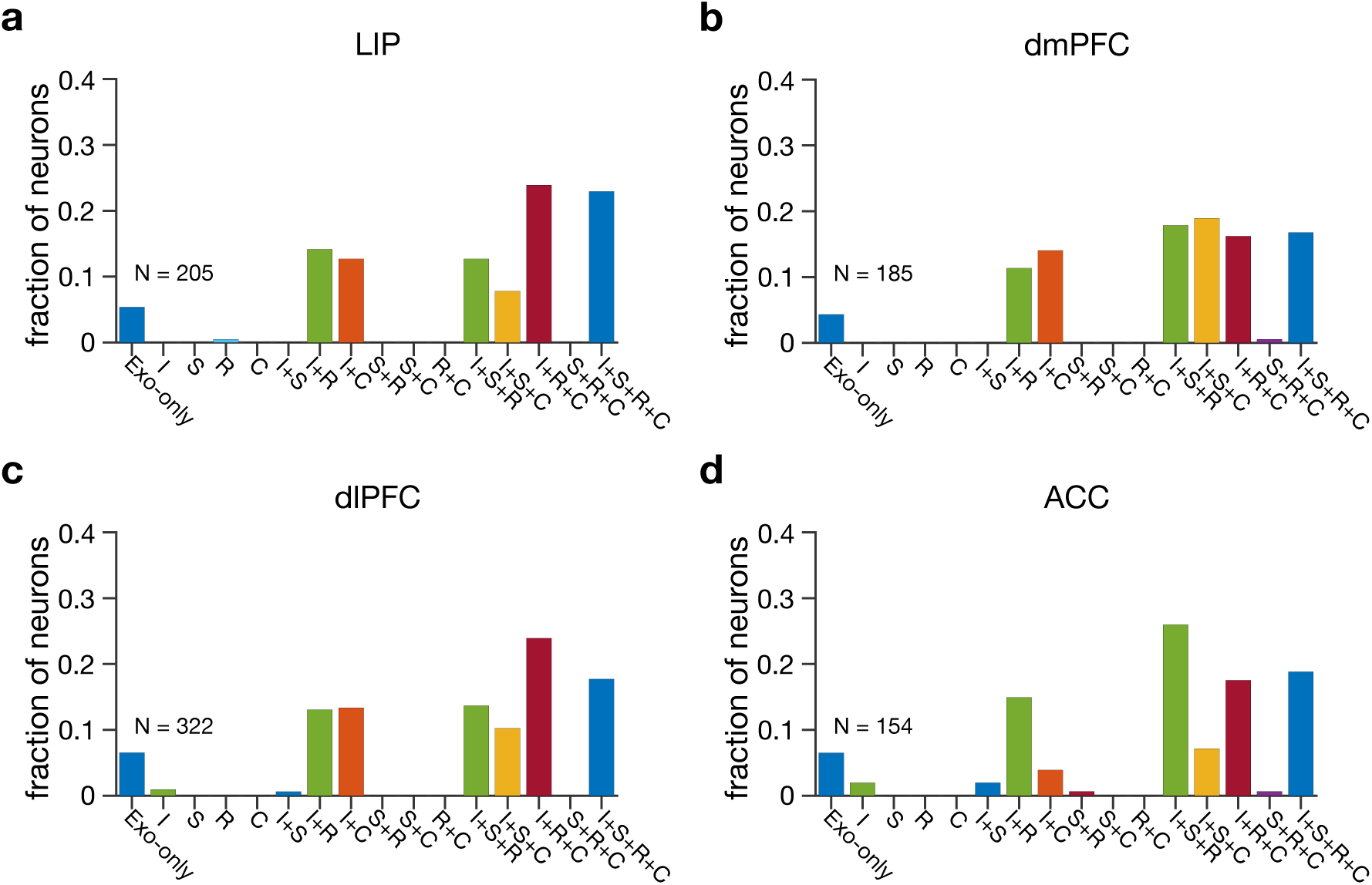
Response of most neurons was best captured using the model with three or more AR or memory components (component with associated timescales). Plots show the fractions of neurons whose activity was best fit by a given model, separately in each of the four cortical areas. Different types of models include: Exo-only, models with only exogenous component and no timescale; I, models with intrinsic timescale; S, models with seasonal timescale; R, models with reward-memory timescale; and C, models with choice-memory timescale. Overall, the models with three or more timescales were able to capture the response of most neurons in all cortical areas (LIP: *fraction*_≥3_ = 0.67, *χ*^2^ = 15.42, *p* = 5.46 × 10^−5^; dmPFC: *fraction*_≥3_ = 0.70, *χ*^2^ = 21.85, *p* = 3.98 × 10^−6^; dlPFC: *fraction*_≥3_ = 0.66, *χ*^2^ = 10.32, *p* = 4.28 × 10^−4^; ACC: *fraction*_≥3_ = 0.70, *χ*^2^ = 12.70, *p* = 7.37 × 10^−4^).

**Supplementary Figure 3.**
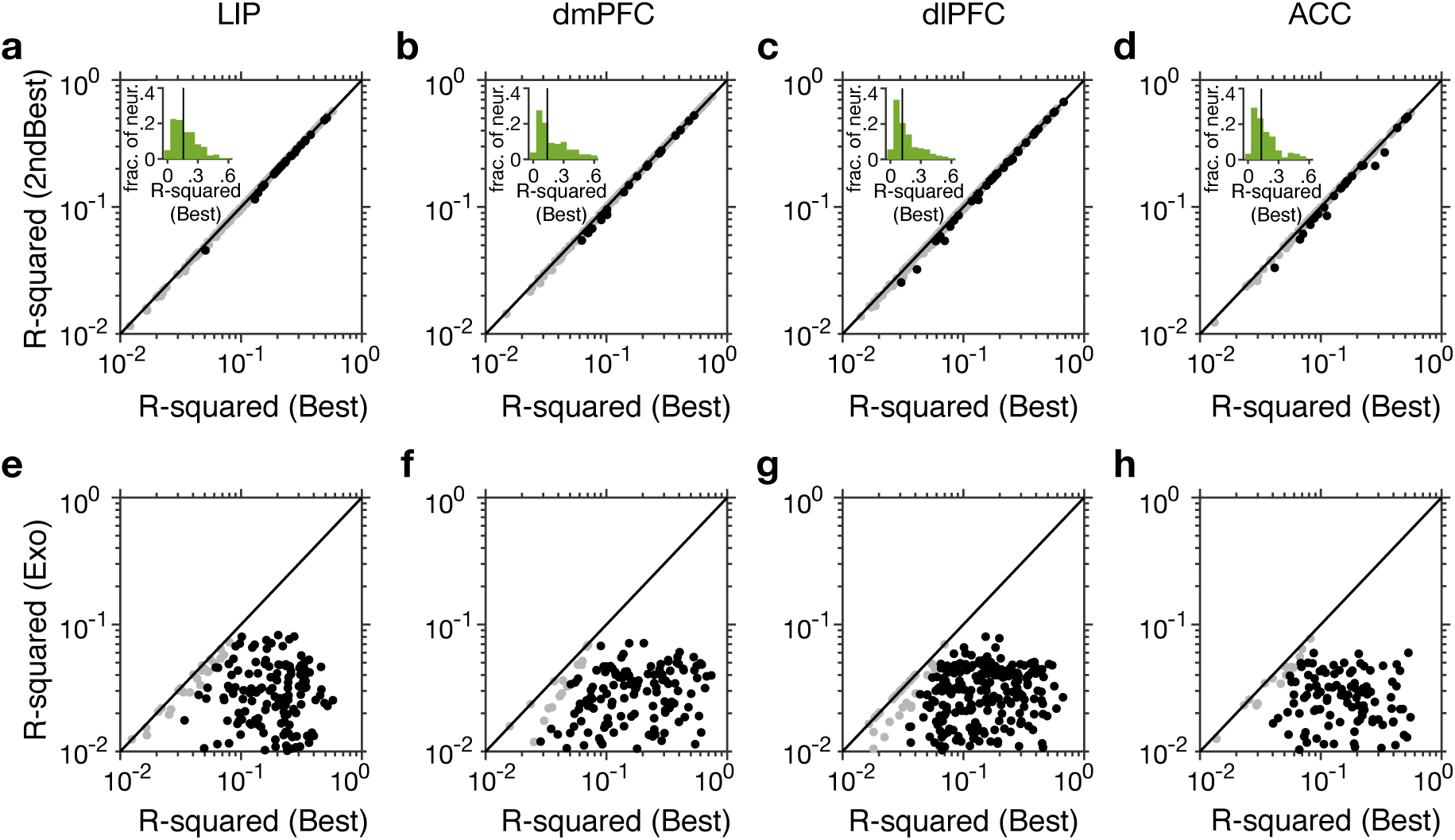
Best model for individual neurons can predict neural response significantly better than the second-best models and models with exogenous component only. Plots show R-squared values for the second-best model vs. those of the best models (**a**–**d**) and R-squared values for the model with exogenous component only vs. those of the best models (**e**–**h**), separately for different cortical areas indicated on the top. Neurons for which the goodness-of-fit for the best model was significantly better than that of the second-based model (or the model with exogenous component only) are shown in black. The insets in the top row show the distributions of the R-squared values for the best models. By definition, R-squared values and the difference in R-squared between the best and second-best models are positive. The differences between the medians of R-squared values for the best model and the model with exogenous component only were significantly different form zero in all cortical areas (two-sided Wilcoxon ranksum test; LIP: *p* = 0.002; dmPFC: *p* = 0.002; dlPFC: *p* = 0.003; ACC: *p* = 0.001).

**Supplementary Figure 4.**
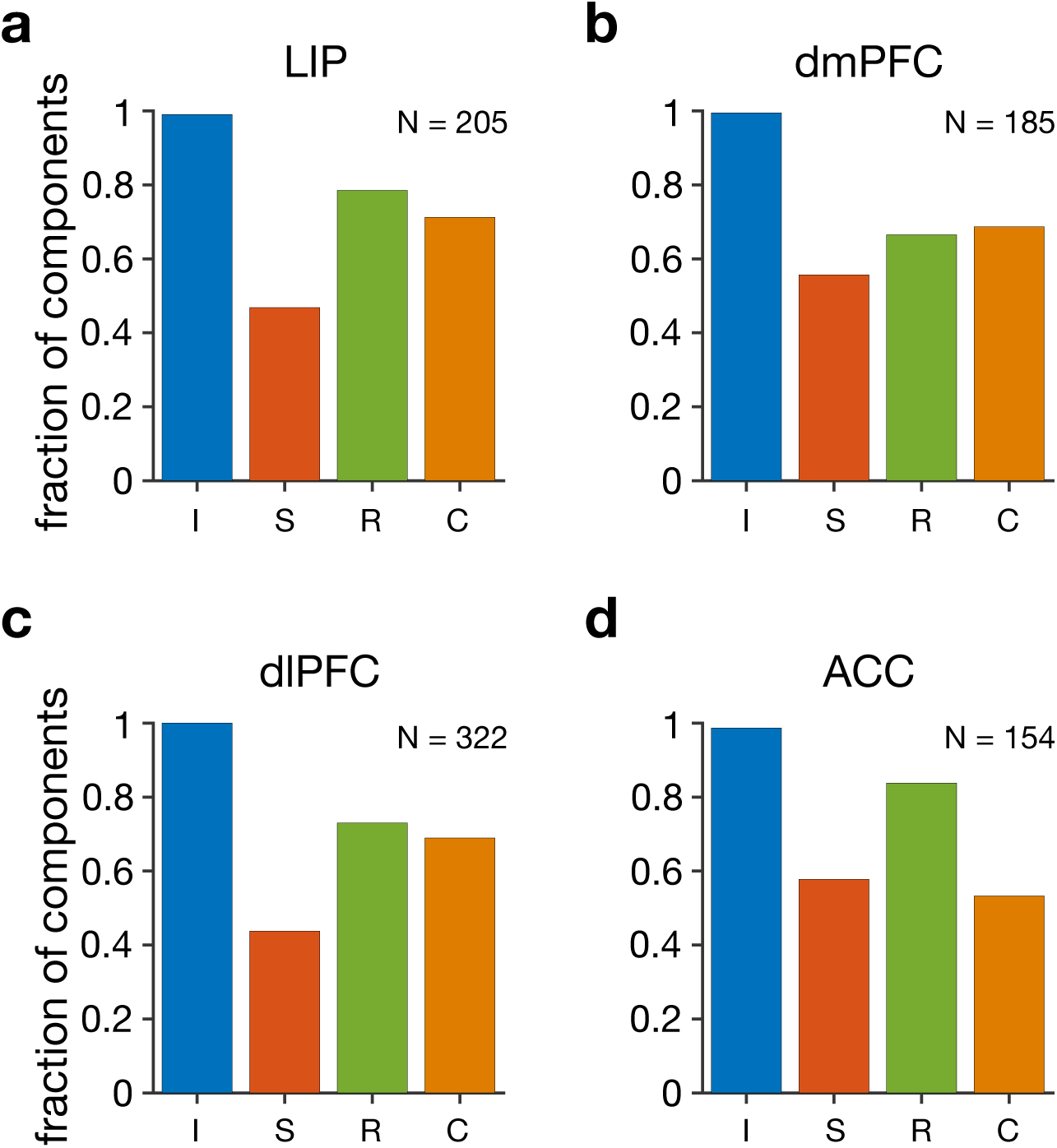
The best model for capturing neural response includes the intrinsic AR component in almost all neurons. Plots show the fractions of different components in the best model across all neurons (I: intrinsic autoregressive; S: seasonal autoregressive; R: reward-memory; C: choice-memory), separately for each cortical area. Reward- and choice-memory components were the second and third most prevalent components of the best models. Nonetheless, the best model of about 50% of neurons also included a seasonal AR component.

**Supplementary Figure 5.**
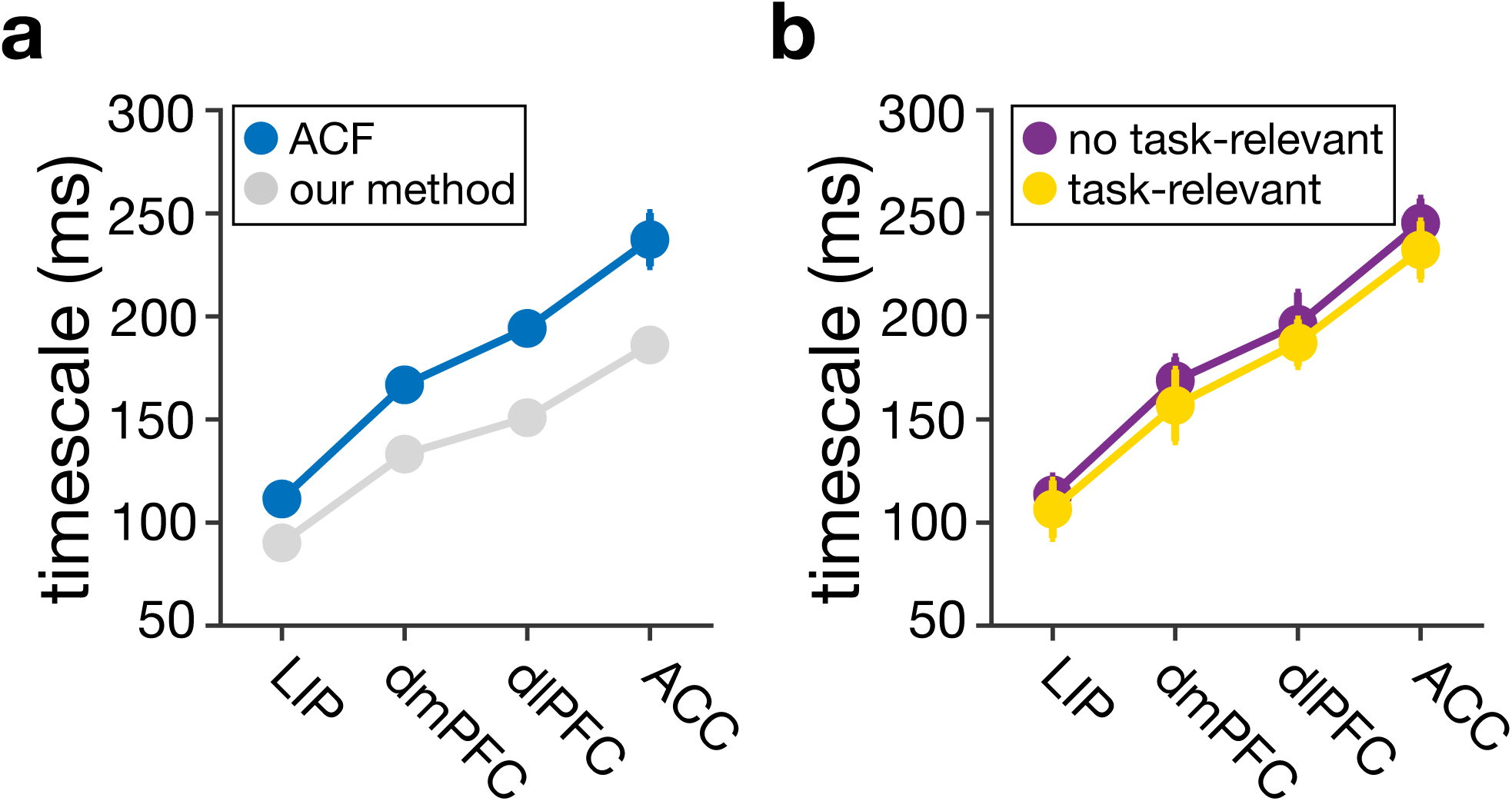
Hierarchy of intrinsic timescales across cortex using activity during the fixation period based on autocorrelation, and lack of evidence for the dependence of intrinsic timescales on the overall selectivity to task-relevant signals. (**a**) Plot (blue symbols) shows the medians of the estimated intrinsic timescales in four cortical areas using autocorrelation function (ACF). Error bars indicate s.e.m. For comparison, the gray symbols show the medians of intrinsic timescales estimated by our method using activity during the fixation period. (**b**) No evidence for the dependence of intrinsic timescales on the overall selectivity to task-relevant (reward outcome and choice) signals based on autocorrelation. Plot shows the median of the estimated intrinsic timescales based on autocorrelation in four cortical areas, separately for neurons with (gold) and without (purple) any type of selectivity to task-relevant signals. There was no significant difference between intrinsic timescales of neurons with and without any task-relevant signals.

**Supplementary Figure 6.**
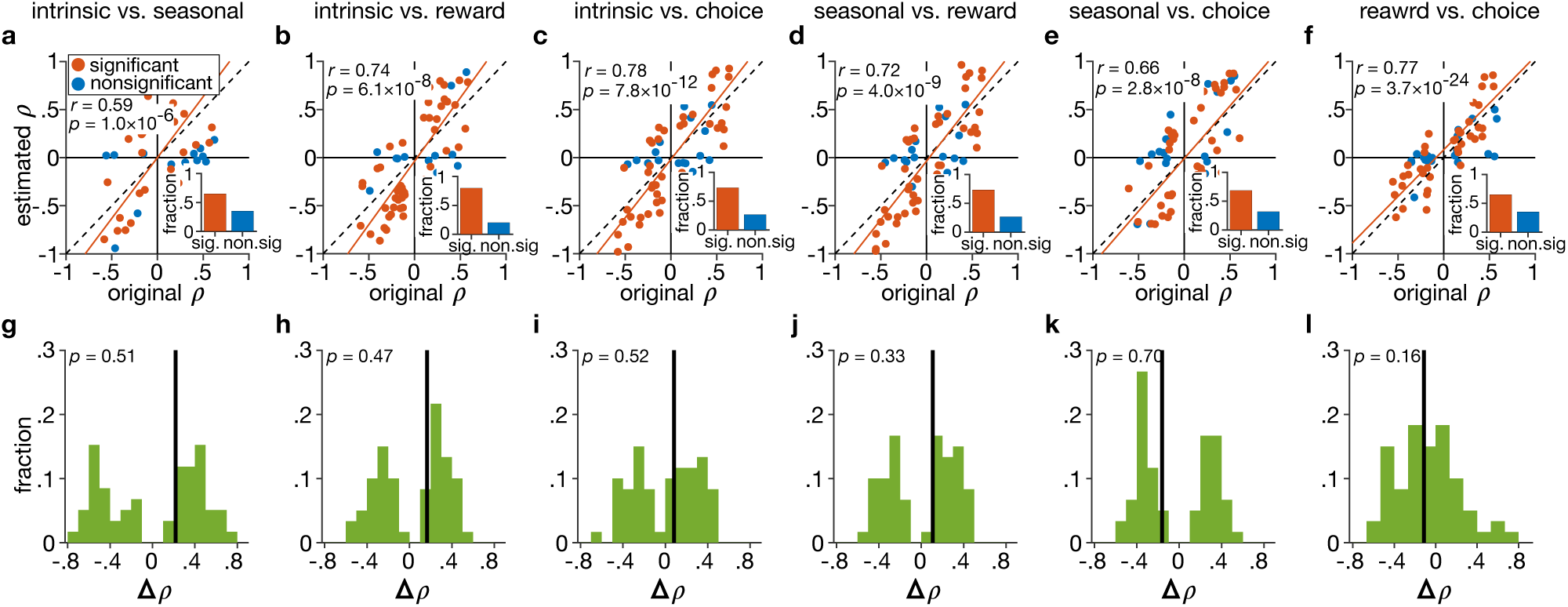
Our method can recover existing correlations between pairs of timescales without any systematic bias. (**a–f**) Plots show estimated vs actual correlations coefficients between a pair of timescales (indicated on the top) across 100 individual neurons in 60 simulated datasets (*N* = 60). Red and blue dots correspond to significant (*p* < 0.05) and non-significant estimated correlations coefficients, respectively (all actual correlation coefficients are significant by design). Insets show the fractions of significant and non-significant estimated correlations. (**g–l**) Plots show the distributions of the difference between actual and estimated correlation coefficients for each pair of timescales indicated on the top. Solid lines indicate the median of each distribution and reported *p*-values are for the test of median being different from 0 (two-sided Wilcoxon ranksum test). The medians of error in estimated correlation coefficients are not significantly different from zero for any pairs of correlation (*p* < 0.05), indicating no bias.

**Supplementary Figure 7.**
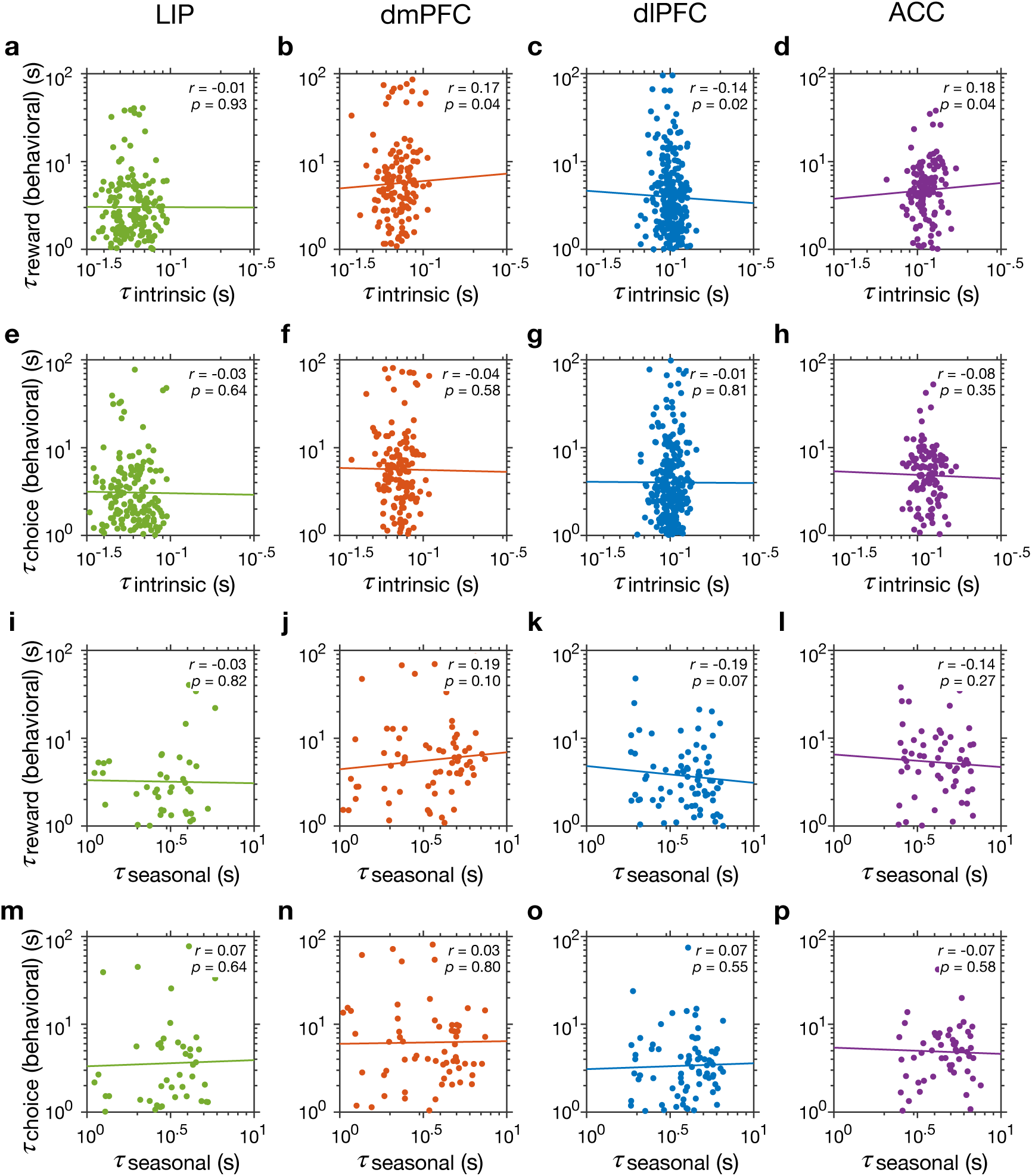
Lack of relationship between the two behavioral timescales and intrinsic and seasonal timescales. (**a–d**) Plots show behavioral reward timescales vs intrinsic timescales of individual neurons recorded during the same sessions, separately for different cortical areas as indicated on the top. Reported are the Spearman correlation coefficients and corresponding *p*-values and the solid lines represent the regression line that was fit to log values. (**e–h**) The same as in a–d but show behavioral choice timescales vs intrinsic timescales. (**i–p**) The same as in a–h but show behavioral timescales vs seasonal timescales. There was no significant correlation between behavioral and neural timescales in any cortical area 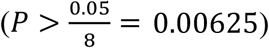.

**Supplementary Figure 8.**
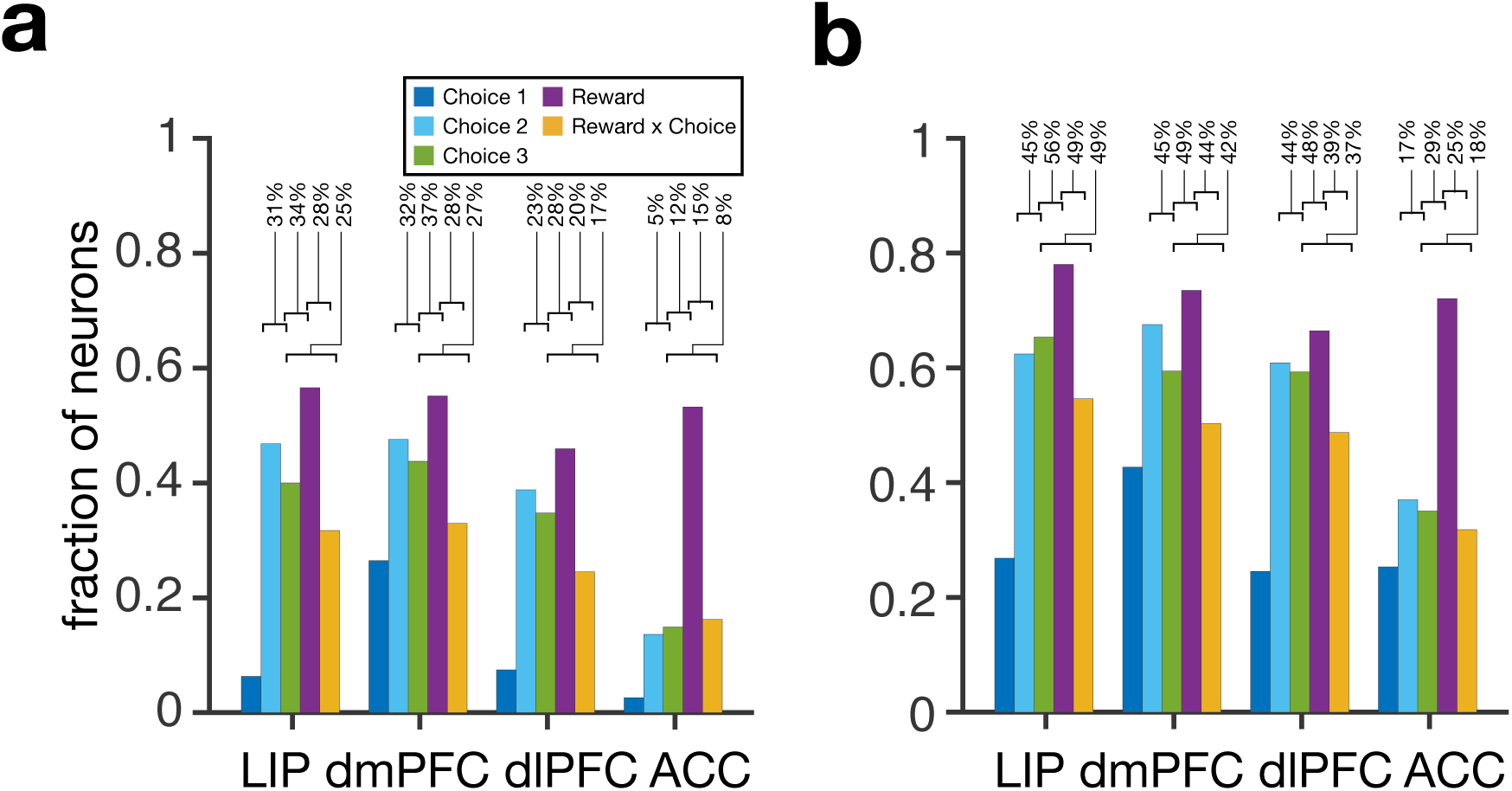
Similar fractions of neurons encode reward outcome across the four cortical areas, whereas the fraction of neurons selective to choice decreases from LIP to ACC. (**a**) Fraction of neurons with selectivity to reward outcome and to choice (in different epochs of the task as indicated in the legend) across the four cortical areas, estimated using the best model for individual neurons. Values on the top indicate the percentage of neurons that exhibit a combination of selectivity in a given area. Overall, about half of neurons in all cortical areas encode reward outcome with no evidence for change in the fraction of reward-selective neurons across cortex (*χ*^2^(3) = 3.49, *p* = 0.062). In contrast, fraction of neurons with choice selectivity decreases from LIP to ACC (*χ*^2^(3) = 23.6, *p* = 1.2 × 10^−6^ for Choice 3). ACC exhibited the smallest fraction of choice-selective neurons (LIP vs. ACC, *p* = 1.3 × 10^−5^; dmPFC vs. ACC, *p* = 1.8 × 10^−6^; dlPFC vs. ACC, *p* = 1.4 × 10^−4^). (**b**) The same as in panel a but using the model that only includes the exogenous term and no timescales.

**Supplementary Figure 9.**
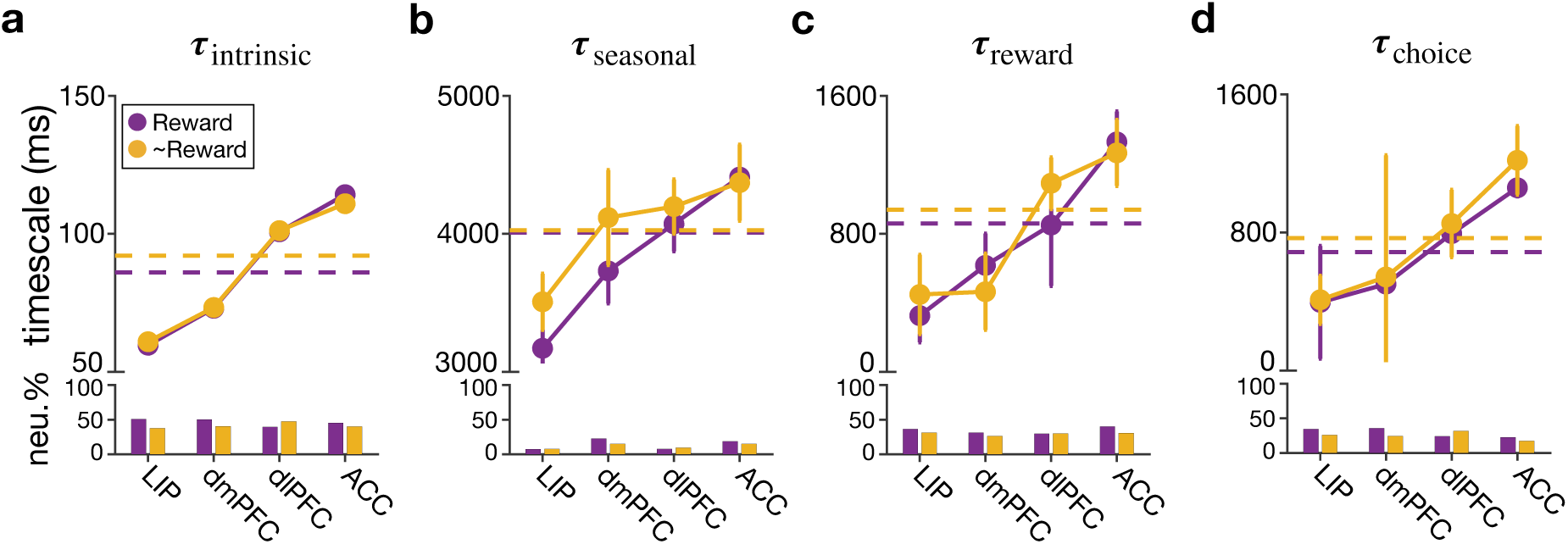
Independence between timescales and the selectivity to reward vs. other task-relevant (choice and interaction of choice and reward) signals. Plots show the median of the estimated intrinsic (**a**), seasonal (**b**), reward-memory (**c**), and choice-memory **(d)** timescales in four cortical areas, separately for neurons selective to reward outcome (purple) and neurons not selective to reward outcome (i.e., those selective to choice or interaction of reward and choice; gold). The dashed lines show the median across all four areas. Error bars indicate s.e.m., and asterisks mark a significant difference between the medians of two types of neurons in a given area or across all areas (two-sided Wilcoxon ranksum test, 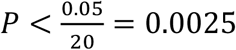 see **Supplementary Table 3** for detailed statistics). Bar graphs show the fractions of neurons selective to reward and neurons selective to non-reward signals in each area. In this analysis, we only included neurons that exhibited selectivity to task-relevant signals.

**Supplementary Figure 10.**
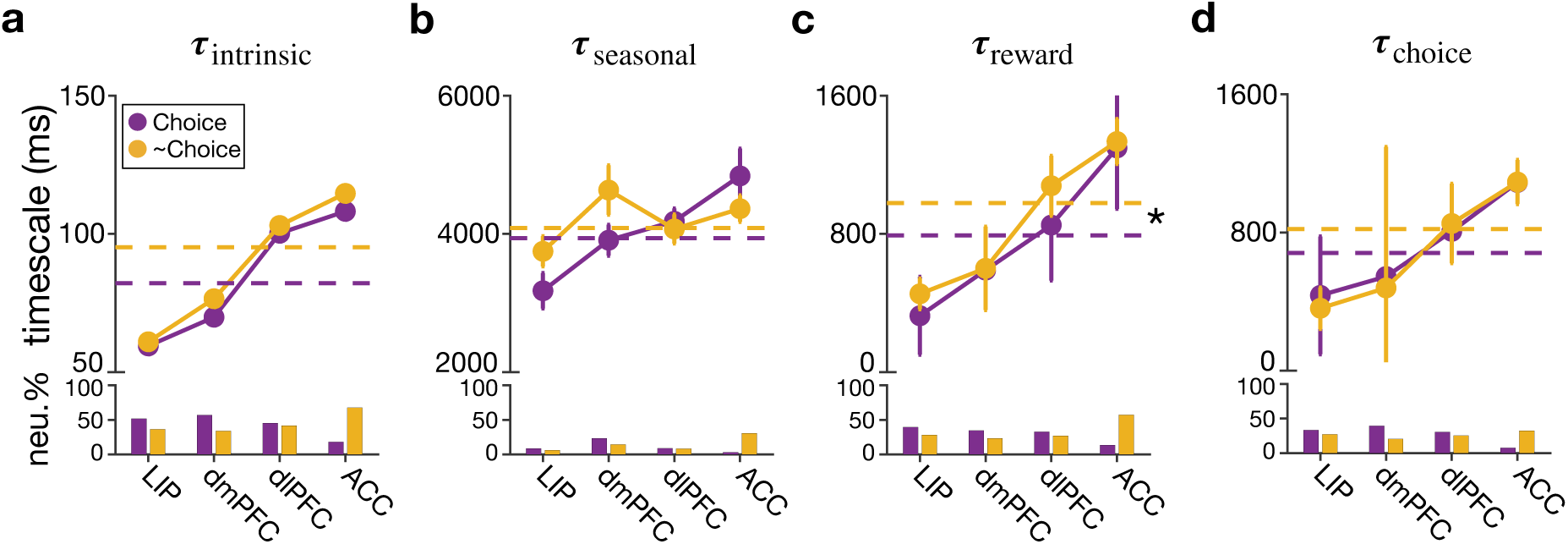
Relationship between timescales and the selectivity to choice vs. other task-relevant (reward and interaction of choice and reward) signals. Plots show the median of the estimated intrinsic (**a**), seasonal (**b**), reward-memory (**c**), and choice-memory (**d**) timescales in four cortical areas, separately for neurons selective to choice (purple) and neurons not selective to choice (i.e., those selective to reward or interaction of reward and choice; gold). The dashed lines show the median across all four areas. Error bars indicate s.e.m., and asterisks mark a significant difference between the medians of two types of neurons in a given area or across all areas (two-sided Wilcoxon ranksum test, 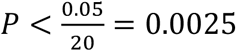 see **Supplementary Table 4** for detailed statistics). Bar graphs show the fractions of neurons selective to choice and neurons selective to non-choice signals in each area. In this analysis, we only included neurons that exhibited selectivity to task-relevant signals.

**Supplementary Figure 11.**
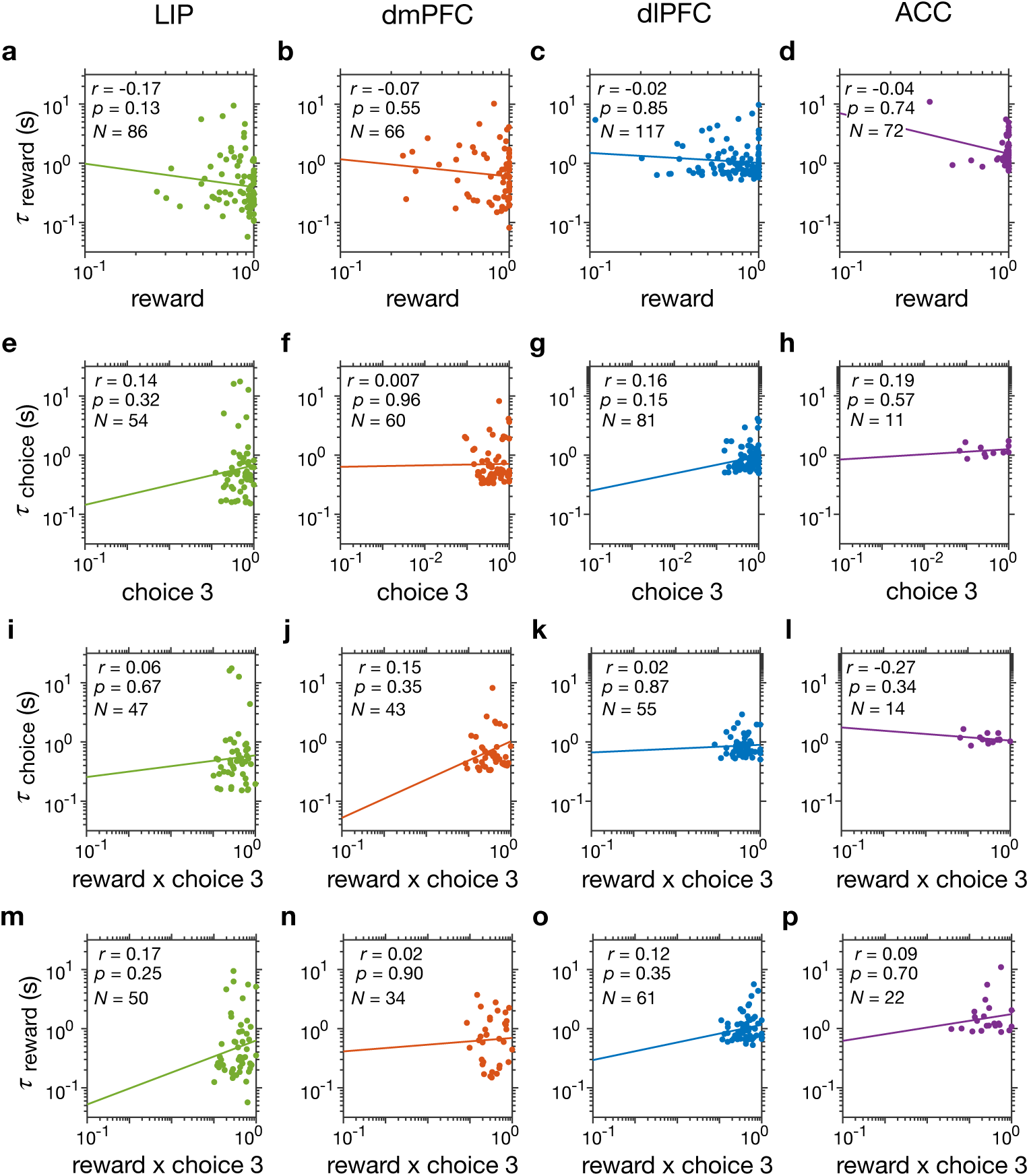
Independence between different types of choice- and reward-memory timescales and magnitudes of response to different task-relevant signals. (**a–d**) Plots show the estimated reward-memory timescales vs. absolute standardized magnitude of reward regressor (as a measure of the selectivity strength) within individual neurons, separately for different cortical areas indicated on the top. Reported are the Spearman correlation coefficients and corresponding *p*-values, and the number of neurons with a significant value of a given timescale. The solid lines represent the regression line that was fit to log values. (**e–h**) The same as in **a–d** but plotting the estimated choice-memory timescales vs. absolute standardized magnitude of choice regressor. (**i–p**) Plots show the estimated choice- (**i–l**) and reward-memory (**m–p**) timescales vs. absolute standardized magnitude of interaction between reward and choice regressors within individual neurons. There was no significant correlation between corresponding memory timescales and exogenous signal magnitude in any of the cortical areas 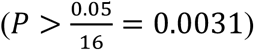.

**Supplementary Figure 12.**
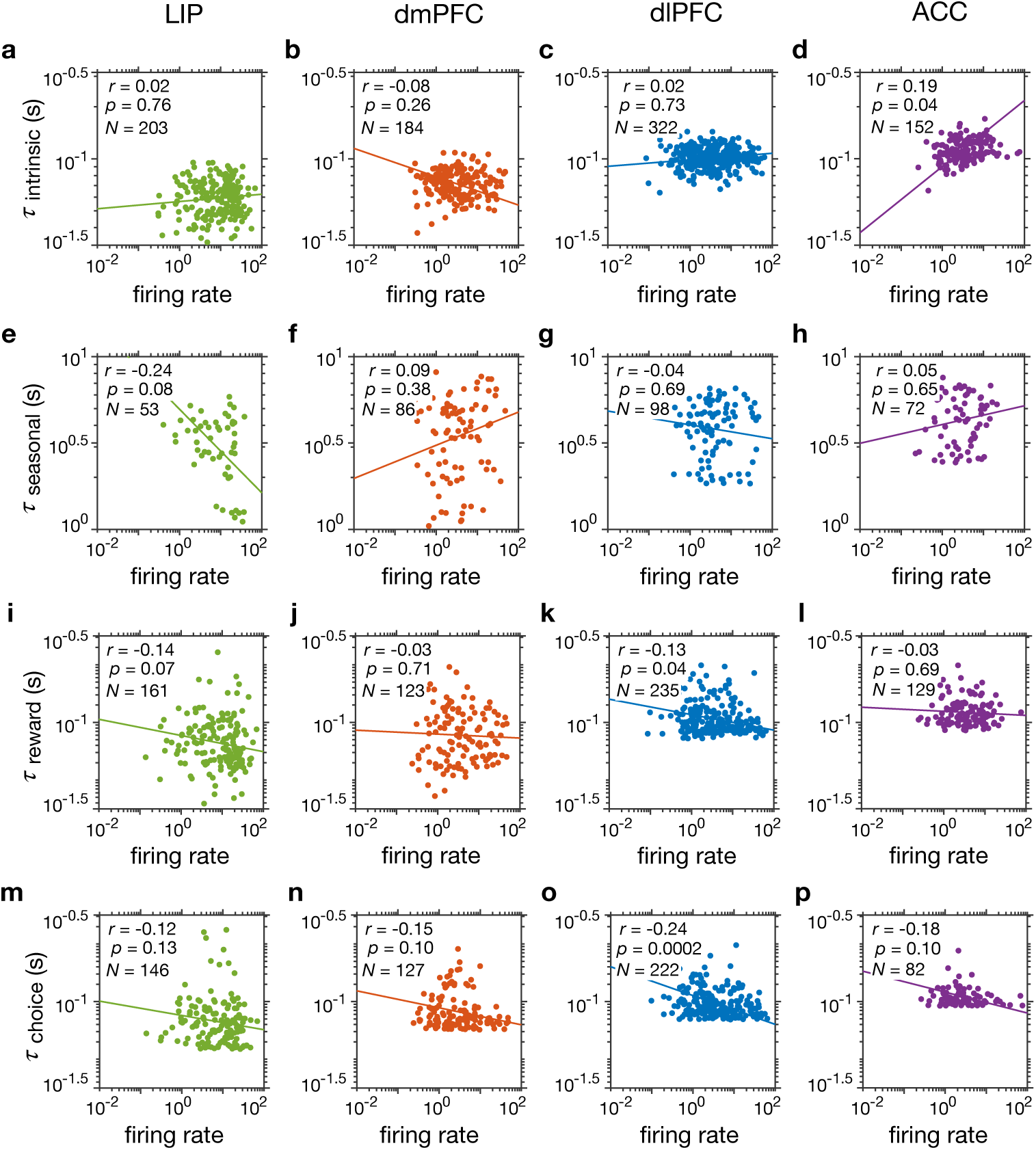
Independence between different types of timescales and firing rates within individual neurons. Panels show scatterplots of estimated intrinsic (**a-d**), seasonal (**e-h**), reward- (**i-l**), and choice- (**m-p**) memory timescales vs. mean firing rates of individual neurons, separately for different cortical areas indicated on the top. Reported are the Spearman correlation coefficients and corresponding *p*-values, and the number of neurons with a significant value of a given timescale. The solid lines represent the regression line that was fit to log values. There was no significant correlation between timescales and firing rates in any of the cortical area 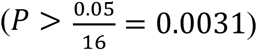.

